# Genetic variability associated with *OAS1* expression in myeloid cells increases the risk of Alzheimer’s disease and severe COVID-19 outcomes

**DOI:** 10.1101/2021.03.16.435702

**Authors:** Naciye Magusali, Andrew C. Graham, Thomas M. Piers, Pantila Panichnantakul, Umran Yaman, Maryam Shoai, Regina H. Reynolds, Juan A. Botia, Keeley J. Brookes, Tamar Guetta-Baranes, Eftychia Bellou, Sevinc Bayram, Dimitra Sokolova, Mina Ryten, Carlo Sala Frigerio, Valentina Escott-Price, Kevin Morgan, Jennifer M. Pocock, John Hardy, Dervis A. Salih

## Abstract

Genome-wide association studies of late-onset Alzheimer’s disease (AD) have highlighted the importance of variants associated with genes expressed by the innate immune system in determining risk for AD. Recently, we and others have shown that genes associated with variants that confer risk for AD are significantly enriched in transcriptional networks expressed by amyloid-responsive microglia. This allowed us to predict new risk genes for AD, including the interferon-responsive oligoadenylate synthetase 1 (*OAS1*). However, the function of *OAS1* within microglia and its genetic pathway are not known. Using genotyping from 1,313 individuals with sporadic AD and 1,234 control individuals, we confirm that the *OAS1* variant, rs1131454, is associated with increased risk for AD and decreased *OAS1* expression. Moreover, we note that the same locus was recently associated with critical illness in response to COVID-19, linking variants that are associated with AD and a severe response to COVID-19. By analysing single-cell RNA-sequencing (scRNA-seq) data of isolated microglia from *APP^NL-G-F^* knock-in and wild-type C57BL/6J mice, we identify a transcriptional network that is significantly upregulated with age and amyloid deposition, and contains the mouse orthologue *Oas1a*, providing evidence that *Oas1a* plays an age-dependent function in the innate immune system. We identify a similar interferon-related transcriptional network containing *OAS1* by analysing scRNA-seq data from human microglia isolated from individuals with AD. Finally, using human iPSC-derived microglial cells (h-iPSC-Mg), we see that *OAS1* is required to limit the pro-inflammatory response of microglia. When stimulated with interferon-gamma (IFN-γ), we note that cells with lower *OAS1* expression show an exaggerated pro-inflammatory response, with increased expression and secretion of TNF-α. Collectively, our data support a link between genetic risk for AD and susceptibility to critical illness with COVID-19 centred on *OAS1* and interferon signalling, a finding with potential implications for future treatments of both AD and COVID-19, and the development of biomarkers to track disease progression.

## Introduction

Alzheimer’s disease (AD) is not only characterised by extracellular amyloid-β deposits, tau tangles and neuronal death, but also extensive neuroinflammatory changes, that may push amyloid pathology to form tau tangles (Edwards, 2019; Ising et al., 2019; Leyns et al., 2019; Yuan et al., 2016). Genetic studies have revealed the importance of gene variants that alter risk for AD and are expressed by the innate immune system, including *APOE*, *TREM2*, *CD33* and *PLCG2* (Efthymiou and Goate, 2017; Hardy and Escott-Price, 2019; Sims et al., 2017). Integrating genetic variants with RNA-sequencing (RNA-seq) and single-cell RNA-sequencing (scRNA-seq) approaches has begun to provide important new insights into the heterogeneous microglial activity changes in AD progression, particularly in the identification of ‘disease-associated microglia’ (DAM) or ‘amyloid-responsive microglia’ (ARM) (Keren-Shaul et al., 2017; Sala Frigerio et al., 2019). This work generally postulates that many AD risk genes, including *APOE*, *TREM2*, *CD33* and *PLCG2,* serve in a related pathway to allow microglia to respond to amyloid-β deposition, altered synaptic activity or damaged phospholipid membranes to activate phagocytosis via the complement system (Edwards, 2019; Hardy and Escott-Price, 2019; Hong et al., 2016; Parhizkar et al., 2019). However, RNA-seq has also identified a parallel trajectory for activated microglia, distinct from DAM/ARM, the so-called ‘interferon-response microglia’ (IRM) (Ellwanger et al., 2021; Friedman et al., 2018; Sala Frigerio et al., 2019; Salih et al., 2019). The number of IRM increases during normal ageing, and increases even further in response to amyloid pathology (Friedman et al., 2018; Sala Frigerio et al., 2019). However, the mechanisms of action of IRM in both settings are not well understood.

Interferons (IFN) are cytokines that trigger a key response to mainly viral pathogens, and consist of three classes: type I (including IFN-α and IFN-β), type II (IFN-γ), and type III (IFN-λ). Interferon signalling is upregulated by an amplification loop in the presence of viruses such as influenza, hepatitis and severe acute respiratory syndrome coronavirus 2 (SARS-CoV-2), leading to the restriction of viral spread, by eliciting a number of molecular effectors to inhibit transcription of viral nucleic acids, digest viral RNA, block translation and modify protein function (Sadler and Williams, 2008). The microglial interferon response has recently been implicated in neuroinflammation and synapse loss in AD, however it is unclear how interferon signalling and anti-viral immune response pathways in microglia contribute to AD pathogenesis (Heuer et al., 2020; Mathys et al., 2019; Olah et al., 2020; Roy et al., 2020; Sala Frigerio et al., 2019; Sebastian Monasor et al., 2020). It has been proposed that nucleic acid-containing amyloid fibrils induce the expression of interferon-stimulated genes in AD-associated mouse models, promoting the upregulation of pro-inflammatory markers and complement cascade-dependent synaptic elimination (Di Domizio et al., 2012; Roy et al., 2020). Another potential contributor to interferon signalling is the DNA-sensing receptor cyclic GMP–AMP synthase (cGAS) and its downstream mediator STimulator of INterferon Genes (STING; cGAS-STING pathway), which respond to not only pathogen-derived nucleic acids but also mitochondrial and genomic nucleic acids derived from stressed, senescent or dying cells in the CNS (Motwani et al., 2019; Paul et al., 2020). While type I interferon signalling was reported to be associated with microglial activation patterns and AD, there are implications that type II interferon signalling also contributes to neuroinflammation and AD pathogenesis (Deczkowska et al., 2016; Majoros et al., 2017; Taylor et al., 2018).

The molecular machinery that drives anti-viral responses includes interferon receptors (IFNAR and IFNGR), tyrosine kinase 2 (TYK2), IFN-stimulated protein of 15 kDa (ISG15, involved in an ubiquitin-like pathway), Mx GTPases (myxovirus resistance), protein kinase R (PKR; also known as eukaryotic translation initiation factor 2-alpha kinase 2, EIF2αK2), and the 2’,5’-oligoadenylate synthetase (OAS)-regulated ribonuclease L. Stimulation of interferon receptors leads to JAK-STAT signalling and induction of expression of interferon-stimulated genes, including *OAS1* from IFN-stimulated response elements (ISREs) in their promoters, resulting in the activation of RNase L which in turn degrades cellular and viral RNA (Donovan et al., 2013; Silverman, 2007). It has recently become apparent that the OAS proteins may also have additional functions that are independent of RNase L activity (Kristiansen et al., 2010; Lee et al., 2019; Li et al., 2016). It has also been shown that *OAS1* is involved in the regulation of cytokine expression (Lee et al., 2019). We recently identified *OAS1* as a putative new risk gene for AD, by integrating DNA sequence variation at the gene-level from human genome-wide association studies (GWAS), associated with AD and a high-resolution RNA-seq transcriptome network expressed by mouse amyloid-responsive microglia (Salih et al., 2019). The amyloid-responsive microglial transcriptome contains several genes involved in interferon signalling, including other *Oas* genes, the interferon receptor *Ifnar2* and the coiled-coil alpha-helical rod protein 1, *Cchcr1*. In parallel work, recent GWAS studies have shown that several genes including *OAS1, IFNAR2* and *CCHCR* involved in interferon signalling also contribute to the genetic risk associated with critical outcomes of COVID-19 and requirement for intensive care (Pairo-Castineira et al., 2021; Schmiedel et al., 2020). Deleterious gene variants have also been described in a number of genes involved in interferon signalling, including *IFNAR1*, *IFNAR2* and *IRF7* in patients with life-threatening COVID-19 pneumonia in an independent study (Zhang et al., 2020). Indeed, neutralising autoantibodies against interferons have been identified in individuals with life-threatening COVID-19, where these antibodies dampen the interferon response (Bastard et al., 2020). These findings implicate the significance of interferon signalling pathways that could exacerbate the progression of AD and the severity of COVID-19.

Similar to neurological manifestations caused by other respiratory viruses including influenza virus and human respiratory syncytial virus (Bohmwald et al., 2018), the latest reports state that as many as 36-78% of people hospitalised with COVID-19 display neurological symptoms including encephalopathy, acute ischaemic cerebrovascular syndrome and neuropsychiatric manifestations (Mao et al., 2020; Paterson et al., 2020; Romero-Sánchez et al., 2020; Woo et al., 2020). Although it is controversial whether SARS-CoV-2 is detectable in the CSF, the virus uses the angiotensin converting enzyme 2 (ACE2) as a key receptor for entry that is expressed by the endothelium of the blood-brain barrier (BBB), epithelial cells of the choroid plexus, and at a lower level by astrocytes, oligodendrocytes and nerve terminals (such as those of the olfactory system, which may allow retrograde spread) (Butowt and von Bartheld, 2020; Chen et al., 2020b; Desforges et al., 2014; Paniz-Mondolfi et al., 2020). There is accumulating evidence that these brain structures and cells are infected with SARS-CoV-2 and are altering the cytokine profile of the brain (Matschke et al., 2020; Yang et al., 2020). Elevated levels of pro-inflammatory cytokines including tumour necrosis factor-α (TNF-α), interleukin (IL)-1 and IL-6 in severe COVID-19 patients (Chen et al., 2020a; Chevrier et al., 2021; Hadjadj et al., 2020; Huang et al., 2020; Keddie et al., 2020; Lee et al., 2020; Qin et al., 2020), trigger an inflammatory cascade leading to cell death. Consequently, a blunted response by the interferon signalling pathway against pathogens implicates a higher risk for developing AD and severe COVID-19. Thus, it is clear that investigating interferon pathways involved in anti-viral responses will: (i) benefit our understanding of pathways involved in the progression of AD and infectious diseases like COVID-19, and (ii) may provide new information for therapeutic approaches and identification of biomarkers to follow disease progression.

In this study, our genotyping analysis confirms that the single nucleotide polymorphism (SNP) rs1131454 within *OAS1* is significantly associated with AD. We find that SNPs within *OAS1* associated with AD also show linkage disequilibrium (LD) with SNP variants associated with critical illness in COVID-19. Additionally, investigating the transcriptome expressed by microglia, we find a genetic co-expression network consisting of genes in interferon response pathways including *OAS1* and the mouse orthologue *Oas1a* are expressed in ageing mice, in mice with amyloid pathology, and in humans with AD and mild cognitive impairment (MCI). Finally, we see that reducing the expression of *OAS1* using siRNA in human induced pluripotent stem cell (iPSC) cells differentiated to microglia, results in an exaggerated pro-inflammatory response when stimulated with IFN-γ. Understanding the mechanisms by which OAS1 pathways bridge interferon and pro-inflammatory signalling in innate immune cells will be important to gain new insights into the progression of AD and severe/critical outcomes associated with COVID-19, and how to treat and track these diseases.

## Materials and Methods

### Genotyping

DNA samples (n=2,547) were kindly obtained from six research institutes in the United Kingdom that are part of the Alzheimer’s Research UK (ARUK) Network. The ARUK series of samples included here are from Bristol, Leeds, Manchester, Nottingham, Oxford and Southampton (further details are given in the Acknowledgements). This ARUK series has not been included in the IGAP consortium study and consisted of confirmed or probable AD diagnosis (n=1,313) and controls (n=1,234). Demographics for these samples are in Table 1. Since all demographic variables were found to be significantly different between the cases and controls (p<0.0001), they were included as covariates in the subsequent analyses. The effects of covariates between cases and controls were tested for age (using a Student’s t-test), and *APOE* status and sex (using chi-squared tests). Samples were genotyped in-house using a TaqMan genotyping assay for rs1131454 following standard protocols (Applied Biosystems); the genotyping rate for the samples was 69.7%. Association of the SNP with AD was conducted using logistic regression correcting for the covariates with the statistical analysis program PLINK (Purcell et al., 2007).

**Table 1.** SNP rs1131454 within OAS1 is associated with Alzheimer’s disease.

| | Mean AAD*<br>(SD) | % Female* | % APOE*<br>E4+ | G-allele<br>frequency | OR G-allele | p value<sup>\$</sup> |
| --- | --- | --- | --- | --- | --- | --- |
| AD<br>(n=1,313) | 76.7<br>(±20.3) | 60.9 | 59.9 | 0.407 | 0.82 | 0.0082 |
| Control<br>(n=1,234) | 72.6<br>(±11.4) | 53.0 | 26.3 | 0.448 |  |  |

Sample demographic and association result for the genotyping carried out on SNP rs1131454 within the *OAS1* gene using the IGAP independent samples from the ARUK DNA bank.
\*Covariates for AD including age at death (AAD), sex and presence of *APOE* E4 alleles were found to be significantly different between cases and controls ( $P < 0.0001$ ). <sup>\$</sup>The association of the G allele for a protective effect was significant after controlling for covariates.

SNPs in the locus containing *OAS1* associated with AD (Salih et al., 2018, 2019), and COVID (Pairo-Castineira et al., 2021), were illustrated with LocusZoom (Pruim et al., 2010). Allele frequencies and LD between these SNPs were investigated using the 1000Genomes Phase 3 data (European populations, CEU and GBR; Auton et al., 2015), and LDlink (Machiela and Chanock, 2015; Myers et al., 2020), where the LD between SNPs was calculated with the LDpair tool (available at https://ldlink.nci.nih.gov/?tab=ldpair using data from European populations, CEU, TSI and GBR).

### RNA-seq data pre-processing

The scRNA-seq datasets were generated from microglia isolated from: i) hippocampi of 3, 6, 12, and 21 month old *APP^NL-G-F^* and WT mice (Sala Frigerio et al., (2019), denoted SF data; non-hippocampal samples were removed), and, ii) dorsolateral prefrontal cortex of human individuals with AD or MCI (Olah et al., (2020), denoted OL data; epilepsy samples and clusters of cells identified by the authors to express high levels of non-microglial or dissociation-induced genes were removed). Data were filtered to remove cells which appeared either unhealthy (>5% ratio of mitochondrial to total unique molecular identifiers (UMIs) for both datasets; <30,000 UMIs or <1,000 genes detected for SF data; <1,000 UMIs or < 700 genes detected for OL data), or potential doublets (>1,000,000 UMIs for SF data; >100,000 UMIs for OL data) using Seurat’s subset function (Stuart et al., 2019). Counts were normalised using Seurat’s *SCTransform* function with the method set to *glmGamPoi*, while regressing out batch effects between sequencing plates with the *vars.to.regress* argument (Hafemeister and Satija, 2019), returning both log-normalised corrected counts and corrected Pearson’s residuals. Genes not reliably detected were removed (zero log-normalised corrected counts in >95% of cells in every cluster of cells assigned by the original authors). Finally, genes showing low variation in expression between cells (coefficient of variation for Pearson’s residuals <15%) were removed, as they are not informative for co-expression analysis.

Bulk RNA-seq datasets generated from microglia isolated from aged and adult wild-type mice (O’Neil et al., 2018), or wild-type and *PSEN2/APP* mice (PS2APP line; Friedman et al., 2018), were normalised using DeSeq2’s *estimatesizefactors* function and transformed by log2(normalised counts+1) (Love et al., 2014). Genes with an average detection level <1.5 normalised counts were removed.

### Co-expression network analysis

Co-expression analysis was performed on pre-processed Pearson’s residuals of the scRNA-seq data from Sala Frigerio et al. (2019), and Olah et al. (2020), using CoExpNets’ *getDownstreamNetwork* function. CoExpNets is an optimisation of the popular weighted gene co-expression network analysis (WGCNA) package (Langfelder and Horvath, 2008), which uses an additional k-means clustering step to reassign genes to more appropriate modules, producing more biologically relevant and reproducible modules (Botía et al., 2017). The expression of the interferon-response modules was represented by averaging expression of the 60 most central module genes in microglia isolated from aged versus young adult wild-type mice, and *PSEN2/APP* versus wild-type mice, and by applying Student’s t-test (the 60 most central module genes were ranked by module membership scores calculated by CoExpNets’ *getMM* function) per sample in processed O’Neil et al. (2018) and Friedman et al. (2018) datasets, respectively.

### Pseudotime analysis

To infer Pseudotime trajectories of microglial activation, we followed the standard workflow of the Monocle 2 R package (Qiu et al., 2017; Trapnell et al., 2014), to calculate single-cell trajectories by ordering cells by their expression of the 1,000 genes most significantly differentially expressed between cell clusters (ranked by qvalue), previously identified by Sala Frigerio et al. (2019).

### Cell culture

Human iPSC-derived microglia cells (h-iPSC-Mg) were generated from the BIONi010-C human iPSC line (EBiSC), originating from a non-demented, normal male (15-19 years old). Differentiation to h-iPSC-Mg was essentially as described by Xiang et al. (2018). Briefly, the protocol was as follows: embryoid body differentiation media for 2 days (consisted of Essential 8, 50 ng/ml BMP-4, 50 ng/ml VEGF, 20 ng/ml SCF, and 10 µM Y-27632), then myeloid differentiation media for 30 days (consisted of X-VIVO 15 medium (Lonza), Glutamax (Life Technologies), 100 U/ml Penicillin/Streptomycin (Life Technologies), 50 µM β-mercaptoethanol (Life Technologies), 100 ng/ml MCSF (Peprotech), and 25 ng/ml IL-3 (Cell Guidance Systems)). These myeloid cells were seeded at a density of 5×10^5^ cells/well in 6-well plates, followed by microglial differentiation media for 13 days (consisted of: DMEM/F12 HEPES no phenol red, 2% ITS-G (Life Technologies), 1% N2 supplement (Life Technologies), 200 µM monothioglycerol (Sigma), Glutamax, NEAA (Life Technologies), 5 µg/ml Insulin (Sigma), 100 ng/ml IL-34 (Peprotech), 25 ng/ml MCSF and 5 ng/ml TGFβ-1 (Peprotech)) and finally, microglia maturation media for 4 days (consisted of: microglial differentiation media plus 100 ng/ml CD200 (Generon), and 100 ng/ml CX3CL1 (Peprotech)).

### siRNA transfection

H-iPSC-Mg were treated with 125 μl Opti-MEM containing 3.75 μl lipofectamine RNAiMAX (#13778150, Thermo Fisher Scientific), according to the manufacturer’s instructions, with 12.5 pmol of either non-targeting negative control siRNA showing no homology to the human transcriptome (#4457287, Ambion), or two different siRNAs targeting *OAS1* (siRNA-1 or −2, Ambion). The siRNA-1 and −2 were selected with an efficiency of 70-95% for *OAS1* knockdown in h-iPSC-Mg. *OAS1* siRNA-1: sense 5’ GUCAAGCACUGGUACCAAAtt 3’, anti-sense 5’ UUUGGUACCAGUGCUUGACta 3’. *OAS1* siRNA-2: sense 5’ CCACUUUUCAGGAUCAGUUtt 3’, anti-sense 5’ AACUGAUCCUGAAAAGUGGtg 3’. H-iPSC-Mg were transfected 4 days after the switch to microglial maturation media. Cells were harvested for RNA extraction 24±1 hr after transfection.

### Interferon-γ treatment

H-iPSC-Mg were treated with 33 ng/ml human IFN-γ (#AF-300-02, PeproTech), at 4 hr following siRNA transfection.

### RNA extraction & cDNA synthesis

Prior to lysis, h-iPSC-Mg cells were washed with Dulbecco’s phosphate-buffered saline (DPBS). Total RNA was extracted using the Monarch Total RNA Miniprep Kit (#T2010S, NEB), following the manufacturer’s guidelines. The total RNA quantity and purity (A_260_/A_280_ ratio) were assessed with a NanoDrop spectrophotometer (Thermo Fisher Scientific). For further reduction of contaminating genomic DNA, equal amounts of RNA were treated with DNase I amplification grade (#18068015, Thermo Fisher Scientific) and RNaseOUT recombinant ribonuclease inhibitor (#10777019, Thermo Fisher Scientific). All RNA samples were reverse transcribed using the LunaScript™ RT supermix kit (#E3010L, NEB), according to the manufacturer’s protocol. In parallel, negative controls were generated with the same procedure except the reverse transcriptase was omitted from the master mix (RT-negative control).

### Real-time quantitative PCR (RT-qPCR)

Primers for genes of interest were designed using Primer-BLAST (NCBI) to test their specificity against the whole human transcriptome. Primers were tested in two-steps. The first step was to resolve the products of a standard PCR on a 3% agarose gel in Tris-acetate-EDTA gel stained with SYBR safe (Thermo Fisher Scientific), to assess the correct product size and presence of only one product. The second step was to test the primers with RT-qPCR using SYBR green to check the linearity range of a dilution series of cDNA template, to test primer efficiency (90-105%), and to confirm primer specificity using a melt curve. Primers with good efficiency were selected (Table S1).

Each RT-qPCR reaction had 1.5 µl of cDNA, 0.25 μM forward and reverse primers, 5 µl of Luna® universal qPCR master mix (#M3003L, NEB), and 3 µl of nuclease-free water. The reactions of each sample across the different conditions were loaded in triplicates with a RT-negative control sample. The reactions were run in 384-well plates on a LightCycler® 480 real-time PCR system (Roche) with cycling conditions of 95◦C for 5 min (preincubation), 45 cycles of [95◦C for 10 sec, 60◦C for 30 sec] (amplification), 95◦C for 5 sec, 65◦C for 1 min, raised to 97◦C (ramp rate 0.11◦C per sec) (melting curve). The product specificity of all reactions was confirmed by the melt-curve analysis, where samples gave only a single peak and RT-negative control samples gave no signal. For normalisation of gene expression levels, the stability of internal control reference genes was tested to ensure the reference genes used were stable across siRNA and IFN-γ treatment. The highest stability was seen with the geometric mean of Ct values of *RPS18*, *GAPDH*, and *HPRT1*. Gene expression analyses were performed by following the GeNorm method (Salih et al., 2012; Vandesompele et al., 2002).

### ELISA of TNF-α secreted by h-iPSC-Mg

For ELISA analyses, supernatants from h-iPSC-Mg cultures were collected, centrifuged to remove cell debris and stored at −20°C. TNF-α levels were quantified using the Quantikine human TNF-α ELISA kit (DTA00D, R&D Systems), following the manufacturer’s instructions. In general, equal volumes of cell supernatants, together with the provided buffer, were loaded in duplicates to the ELISA microplate and incubated for 2 hr at RT. Following four washes, wells were then incubated with horseradish peroxidase conjugated human TNF-α antibody for another 2 hr at RT. The microplate was washed again and incubated with a substrate solution prepared with chromogen (TMB) and hydrogen peroxide for 30 min before addition of a stop solution with sulphuric acid. Then the colour intensity was measured at 450 nm, and 540 nm to assess the background signal, using a FLUOstar Omega microplate reader and the TNF-α concentrations in h-iPSC-Mg supernatants were calculated from the standard curve.

### Statistical analyses for cell culture experiments

All statistical analyses were conducted on GraphPad Prism 9. For experiments with h-iPSC-Mg under basal conditions, comparisons between multiple groups were analysed by using repeated-measures one-way ANOVA (repeated for wells treated with different siRNA in the same plate) followed by Dunnett’s multiple comparisons tests comparing every test group with the negative control group. For experiments with h-iPSC-Mg in response to IFN-γ treatment and siRNA treatments, comparisons between factors were tested with two-way ANOVA followed with Tukey’s multiple comparisons tests to compare all possible permutations of sample groups. Significant differences were indicated as follows: *p<0.05, **p<0.01, ***p<0.001, ****p<0.0001. Data are shown as mean ± standard error of the mean (SEM), where the sample size (N) represents individual cell preparations.

## Results

### SNPs associated with OAS1 are linked to risk for AD and critical illness with COVID-19

Our recent work overlapping the mouse amyloid-associated transcriptome network (bulk RNA-seq) with human gene-level-aggregated AD risk variants, identified a microglial co-expression network, whose eigengene strongly correlated with the level of amyloid-β pathology and contained the mouse orthologues of many known human GWAS loci (including *TREM2* and *APOE*) (Salih et al., 2019). Our recent work also predicted the importance of several previously unidentified risk genes, including *OAS1*, and demonstrated a colocalisation between AD risk loci and eQTLs regulating *OAS1* expression in: (i) human iPSC-derived macrophages stimulated with a combination of IFN-γ and salmonella, and (ii) human monocytes stimulated with LPS or 5′-triphosphate double-stranded RNA (Alasoo et al., 2018; Kim-Hellmuth et al., 2017; Salih et al., 2019). To further evaluate AD-associated genetic variation in *OAS1*, we genotyped the top SNP, rs1131454, identified by our gene-level analysis of the Lambert *et al*. GWAS (Lambert et al., 2013). SNP rs1131454 was genotyped in 1,313 individuals with sporadic AD and 1,234 control individuals that were not included in the original cohort from Lambert et al. (2013). We found a significant association of rs1131454 with AD in this independent cohort of AD and control individuals (Table 1).

To determine whether AD-associated SNPs in close proximity to *OAS1* were in LD with recently identified SNPs related to critical outcomes with COVID-19 (Pairo-Castineira et al., 2021), we used the 1000Genomes Phase 3 database (CEU and GBR populations; Auton et al., 2015) (Fig. 1A and 1B). We saw a strong LD (r^2^=0.63-0.99 and D’=0.91-1.0) between our two top SNPs adjacent to *OAS1*, rs1131454 (identified using data from Lambert et al. (2013)), and rs4766676 (identified using data from Kunkle et al. (2019)), and two SNPs associated with severe COVID-19 responses, rs6489867 and rs10735079 (Pairo-Castineira et al., 2021) using LDpair. This suggests that four risk alleles adjacent to the *OAS1* and *OAS3* genes may form a haplotype that contributes to both risk for AD and a severe response with COVID-19. Together with results from our previous AD-associated colocalisation analyses (Salih et al., 2019), the recent transcriptome-wide association study (TWAS) for critically ill COVID-19 patients, suggests that there may be complex or pleiotropic variant(s) within the *OAS* gene cluster that modulates risk for both diseases potentially via regulation of *OAS1* or *OAS3* expression (Pairo-Castineira et al., 2021; Salih et al., 2019; Schmiedel et al., 2020).

**FIGURE 1.**
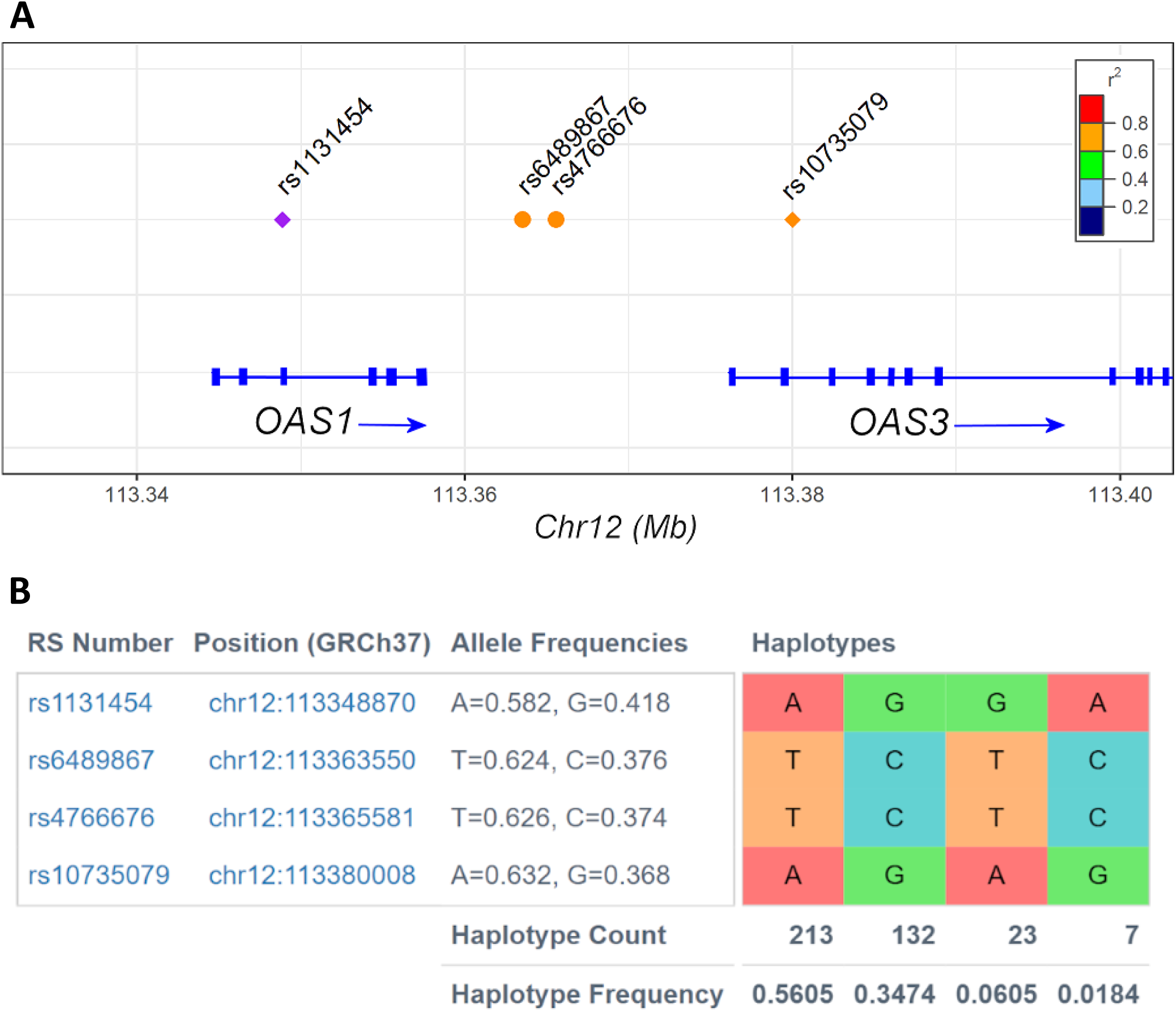
The genomic position of four SNPs on 12q24: rs1131454 and rs4766676 (associated with AD) and rs10735079 and rs6489867 (associated with critical illness with COVID-19) relative to *OAS1*. (A) We have confirmed rs1131454 is associated with AD (Table 1) following our analysis of AD GWAS data (our data is given in Salih et al. (2018), which was derived from further analysis of the GWAS data in Lambert et al. (2013)). We identified the association of rs4766676 with AD (our data is given in Salih et al. (2019), which was derived from further analysis of the GWAS data in Kunkle et al. (2019)). SNPs rs10735079 and rs6489867 are associated with severe illness with COVID-19 (Pairo-Castineira et al., 2020). The R^2^ scale indicates LD between rs1131454 (the reference SNP, purple), compared to the other three SNPs. Drawn with LocusZoom (Pruim et al., 2010). (B) Strong correlation between the SNPs associated with AD and COVID-19 suggesting they might form part of a haplotype group. Allele frequency at these SNPs was investigated using the 1000Genomes Phase 3 data (CEU and GBR populations; Auton et al., 2015) and LDlink (Machiela and Chanock, 2015; Myers et al., 2020).

### Genetic co-expression network involving interferon response genes is increased with ageing and amyloid plaques

To understand how the expression of *OAS1* relates to microglial activation states, we initially assessed the expression of its mouse orthologue, *Oas1a*, in microglia isolated from *APP^NL-G-F^* knock-in and wild-type C57BL/6J mice at 3 to 21 months of age, profiled by scRNA-seq (Sala Frigerio et al., 2019). We identified a co-expression transcriptomic network containing *Oas1a* by utilising a modified version of WGCNA, which produces more biologically relevant co-expression networks (Fig. 2A). The module containing *Oas1a* displayed many interferon-responsive genes (*Ifit1*, *Ifit2*, *Ifit3*, *Ifitm3*, *Stat1*, *Stat2, Usp18* and *Mx1*), as expected, and pro-inflammatory cytokines (*Tnf*) (Table S2). We then generated a two-branched diffusion trajectory by semi-supervised pseudotime analysis, to visualise two different activation states ARM and IRM derived from homeostatic microglia (Fig. 2B). This revealed that *Oas1a* and the interferon response module were upregulated in microglia that transition along one branch of this trajectory towards the IRM state (Fig. 2C). This module showed increased expression in microglia isolated and pooled (bulk RNA-seq) from aged wild-type mice compared to young mice (Fig. 2D). Moreover, this interferon module showed increased expression in microglial isolated and pooled from *PSEN2/APP* mice relative to wild-type mice (Fig. 2E). Importantly, performing similar analyses in microglia isolated from human brains with AD (Olah et al., 2020), we identified a related IRM network containing *OAS1* and other interferon responsive genes (overlap with mouse IRM network p=3.7e-13, Fisher’s Exact test) (Fig. 2F) (Table S3). These findings suggest that a discrete population of interferon-responsive microglia defined by co-expression of *OAS1* and pro-inflammatory cytokines may play a role in amyloidosis, as well as ageing, the major risk factor for AD and COVID-19 (Guerreiro and Bras, 2015; Ou et al., 2020).

**FIGURE 2.**
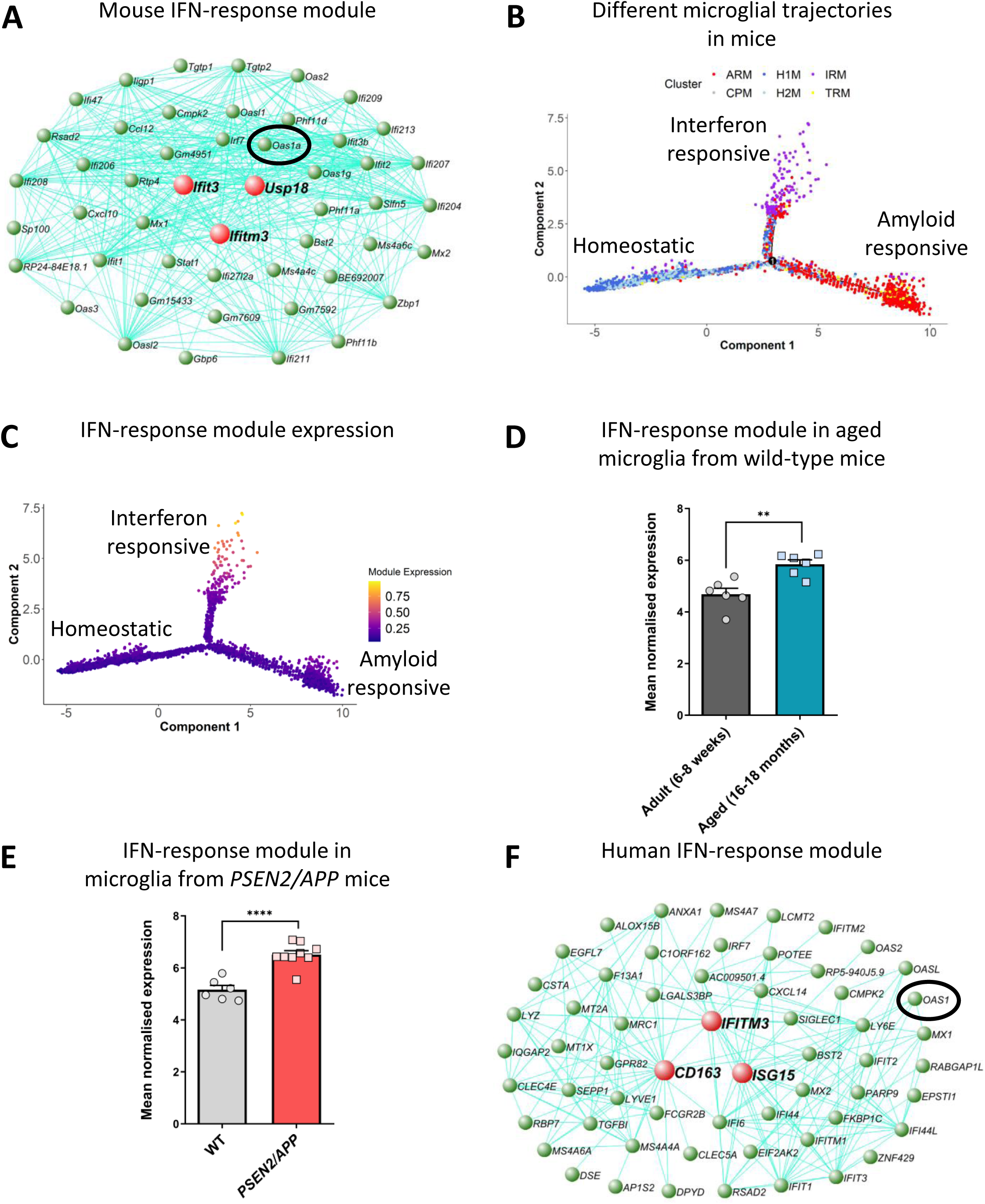
An interferon-response associated gene module is present along a distinct microglial activation trajectory upregulated in aged mice, mice with amyloid pathology and in humans with AD. A) The genetic network containing *Oas1a* from microglial cells isolated from wild-type and *APP^NL-G-F^* KI mice at 3, 6, 12 and 21 months of age analysed by scRNA-seq (Sala Frigerio et al., 2019). The 50 genes showing the highest connectivity are plotted, and *Oas1a* is highlighted. Green nodes represent genes, edge lines represent co-expression connections, and the central large red nodes are the hub genes (full network given in Table S2). (B) Semi-supervised pseudotime ordering of microglial cells isolated from wild-type and *APP^NL-G-F^* KI mice as above based on expression (Sala Frigerio et al., 2019), with Monocle 2, shows homeostatic cells as the root state, and ARM and IRM as the endpoints of distinct activation trajectories. C) The gene module containing *Oas1a* is upregulated along the IRM-associated activation trajectory. The expression of this module is relatively absent from both the root homeostatic state and the ARM trajectory. D) Mean normalised expression of the 60 most central genes in the interferon response module is greater in microglia isolated from aged wild-type relative to young adult mice (6-8-weeks versus 16-18-months of age; N=6 mice per group. Data shown as mean ± SEM. Student’s t-test; **p<0.01. Further analysis of data from O’Neil et al. (2018)). E) Mean normalised expression of the 60 most central genes in the interferon response module is greater in microglia isolated from *PSEN2/APP* relative to wild-type mice at 14-15-months of age (N=6-9 mice per group. Data shown as mean ± SEM. Student’s t-test; ****p<0.0001. Further analysis of data from Friedman et al. (2018)). F) Genetic network plot of an interferon-response associated module detected in microglial cells isolated from human AD patients and individuals with MCI (Olah et al., 2020). This module shows a significant overlap with the interferon-response module detected in mice (see panel A) (full network given in Table S3). The hub gene of this module is *CD163*, and a number of other macrophage marker genes are prominent within this module (*MSR1*, *TGFB1*, *F13A1*, *LY6E* and *LYZ*), indicating that expression of *OAS1* and other interferon-response genes in the human AD patient brain are either associated with a population of CNS border-associated or invading macrophages, or potentially a microglial subpopulation which upregulates anti-inflammatory genes. *OAS1* is highlighted.

### *OAS1* coordinates the pro-inflammatory response of human iPSC derived microglia

To study the function of *OAS1* in human microglia, we used siRNA to knockdown expression of *OAS1* in h-iPSC-Mg. We obtained knockdown of *OAS1* expression to around 30-40% of endogenous levels under non-stimulated conditions (Fig. S1). Under these basal conditions, the expression profiles of commonly used markers covering various microglial functions including homeostasis, phagocytosis, pro- and anti-inflammatory signalling, and genes from the interferon-responsive networks, generally remained unchanged by *OAS1* knockdown in h-iPSC-Mg. In response to IFN-γ treatment, *OAS1* expression increased by around 3.5-fold, and *OAS1* knockdown with siRNA was still effective in the presence of IFN-γ (Fig. 3A). Treatment with IFN-γ also induced the expression of pro-inflammatory marker *TNF* by around 30-fold in h-iPSC-Mg, but with *OAS1* knockdown the induction of *TNF* was further significantly increased to around 45-fold compared to control cells treated with non-targeting siRNA (Fig. 3B). We confirmed the increased secretion of TNF-α into the h-iPSC-Mg culture media in samples with *OAS1* knockdown in the presence of IFN-γ compared to cells treated with non-targeting siRNA and IFN-γ (Fig. 3C). At the same time, *C1QA* expression showed a significant but modest increase with *OAS1* knockdown (Fig. 3D). Treatment with IFN-γ resulted in increased expression of *TGFB1*, *CD68*, *P2RY12*, *IFITM3* and *STAT1*, decreased expression of *TREM2*, and had no effect on *IL1B*, *ITGAM* and *CD163* expression, with *OAS1* knockdown not altering the expression of these genes (Fig. S2). Therefore, *OAS1* appeared to dampen the pro-inflammatory response involving TNF-α following IFN-γ stimulation, with a modest effect on dampening the expression of *C1QA* complement.

**FIGURE 3.**
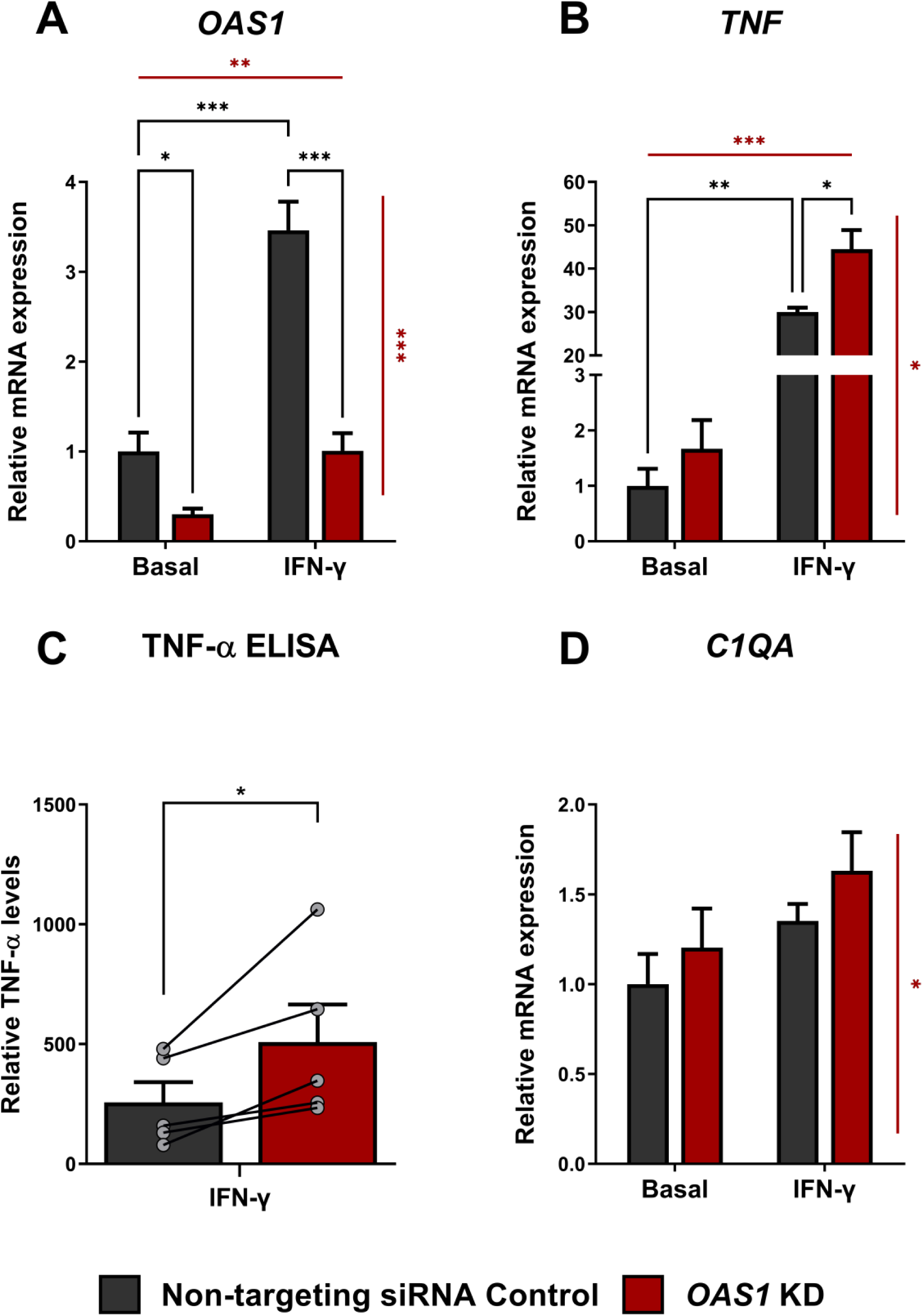
*OAS1* knockdown in the presence of IFN-γ resulted in exaggerated expression of *TNF* in h-iPSC differentiated to microglia. (A) *OAS1* expression was knocked-down using siRNA under basal conditions and in response to IFN-γ. Two-way ANOVA with significant main effects of IFN-γ treatment (p<0.01), and siRNA treatment (p<0.001) indicated as horizontal and vertical red lines, respectively, with a significant interaction (p<0.001). (B) *TNF* expression, as a marker of the pro-inflammatory response, showed significant up-regulation in response to IFN-γ application, and accentuated expression with *OAS1* knockdown. Two-way ANOVA with significant main effects of IFN-γ treatment (p<0.001), and siRNA treatment (p=0.015) indicated as horizontal and vertical red lines, respectively, with a significant interaction (p=0.023). (C) ELISA of TNF-α levels in the conditioned h-iPSC-Mg media. TNF-α levels were normalised to the geometric mean of *GAPDH*, *HPRT1* and *RPS18* assessed by RT-qPCR. Note: TNF-α protein was not detectable in the media of h-iPSC-Mg not treated with IFN-γ. N=5 independent plates. Data shown as mean ± SEM. Paired Student’s t-test (paired for wells on same plate); * p<0.05. (D) *C1QA* expression was increased with *OAS1* knockdown. Two-way ANOVA with significant main effect of only siRNA treatment (p=0.029), not IFN-γ treatment (p>0.05), indicated as a vertical red line, with no significant interaction (p>0.05). Gene expression levels were normalised to the geometric mean of *GAPDH, HPRT1* and *RPS18*, then calculated as fold change relative to the non-targeting siRNA control (vehicle control) without IFN-γ treatment in each individual culture preparation. N=5 independent plates. Data shown as mean ± SEM. Two-way ANOVA; significant main effects of IFN-γ treatment and *OAS1*-knockdown indicated by red lines. When a significant interaction was seen between IFN-γ treatment and siRNA treatment, Tukey’s multiple comparisons tests were then performed to test for pairwise significance of all groups; * p<0.05, ** p<0.01, *** p<0.001, **** p<0.0001.

## Discussion

GWAS studies have revealed the importance of innate immune cells in moderating the risk of AD, with the discovery of variants in genes such as *TREM2*, *CD33* and *PLCG2*. A number of studies have shown that several of these genes are likely involved in the same signalling pathway or functioning in the same sub-population of microglia to control their activation to a so-called DAM or ARM state (Griciuc et al., 2019; Keren-Shaul et al., 2017; Krasemann et al., 2017; Sala Frigerio et al., 2019). These DAM or ARM cells are thought to regulate complement-dependent phagocytosis of synapses, which may be dependent on phospholipid signals presented on damaged cell membranes (Edwards, 2019; Györffy et al., 2018; Hardy and Escott-Price, 2019; Hong et al., 2016; Scott-Hewitt et al., 2020). Emerging data is also indicating that other sub-populations of microglia play a significant role in AD and ageing, particularly the IRM cells, and may also express genes that confer risk for AD (Friedman et al., 2018; Olah et al., 2020; Sala Frigerio et al., 2019; Salih et al., 2019). We recently identified the interferon-responsive gene *OAS1* as a putative risk gene for AD by combining transcriptional changes in the presence of amyloid plaques and gene-level variation from GWAS data (Salih et al., 2019). Here, we confirm that rs1131454 within *OAS1* is associated with AD when we genotype an independent cohort of 1,313 individuals with AD and 1,234 control individuals. Additionally, we show that rs1131454 is in LD with newly identified SNPs associated with critical illness due to COVID-19, suggesting that the same locus regulates the risk for both AD and severe outcomes with COVID-19. Furthermore, by building genetic transcriptome networks using scRNA-seq of isolated mouse microglia, we show that an interferon response pathway containing the mouse orthologue of *Oas1a* exhibits increased expression during ageing in microglia. This indicates that the expression of the interferon pathway occurs within a specific sub-population of innate immune cells and contributes to age-dependent changes that may predispose or protect some people in the population against these age-related diseases. We identify a related network of interferon-responsive genes containing *OAS1* in human microglia isolated post-mortem from individuals with AD. Finally, with functional experiments using h-iPSC-Mg, we show that *OAS1* levels moderate the pro-inflammatory response of myeloid cells in response to elevated interferon levels. Thus, individuals with lower levels of *OAS1* due to eQTL variants can be more likely to show a strong pro-inflammatory response to AD-associated pathology and COVID-19, triggering damage and cell death in neighbouring cells such as neurons and alveolar cells, by initiating a ‘cytokine storm.’

The data presented here provide support for *OAS1* linking AD and critical illness with COVID-19 by controlling the pro-inflammatory output of myeloid cells. OAS1 is an oligoadenylate synthetase enzyme that binds dsRNA and changes conformation to generate oligoadenylates and activate RNase L to digest RNA (Schwartz and Conn, 2019; Schwartz et al., 2020). It has also been proposed that OAS1 may also have RNase-independent roles (Kristiansen et al., 2010; Li et al., 2016). *OAS1* is expressed by multiple cell types in the myeloid lineage including monocytes, macrophages, natural killer cells and microglia (Alasoo et al., 2018; Kim-Hellmuth et al., 2017; Schmiedel et al., 2018, 2020). Here, we show that rs1131454 within *OAS1* is associated with AD, and previously we showed that this locus acts as an eQTL regulating the expression of *OAS1* in monocytes and iPSC-macrophages stimulated with 5’-triphosphate dsRNA and a combination of IFN-γ and salmonella, respectively (Salih et al., 2019). Also in that locus is *OAS2*, which was independently shown to be associated with AD (Broce et al., 2019; Kuksa et al., 2020). Recent work has identified other SNPs close to this locus associated with severe responses to COVID-19 that also act as eQTLs and regulate the expression of *OAS1*, *OAS3* and other distal genes such as *DTX1* (Pairo-Castineira et al., 2021; Schmiedel et al., 2020). Indeed, a series of different variants have been identified within *OAS1* that modify susceptibility to infection with SARS-CoV-2, other viruses such as hepatitis C, developing type 1 diabetes, and multiple sclerosis; these variants have been proposed to cause amino acid changes, altered splicing (spliceQTLs) and altered expression (eQTLs) (Bonnevie-Nielsen et al., 2005; Cagliani et al., 2012; He et al., 2006; Klaassen et al., 2020; Randolph et al., 2020; Tessier et al., 2006; Zeberg and Pääbo, 2021). A recent study has identified a haplotype of approximately 75 kb centred around the *OAS* locus inherited from Neandertals that is protective against COVID-19 (Zeberg and Pääbo, 2021), suggesting that there has been genetic selection at this locus during European history. Given the variety of SNPs associated with *OAS1* and their influence on diseases related to innate immune function, this suggests that these SNPs tag a complex and/or pleiotropic genetic variant that alters the expression of *OAS1*, as opposed to a variant that alters *OAS1* enzyme activity. The different SNPs tagging the *OAS1* variant may be in different chromosomal positions in different people, with different effect sizes. It is also possible that varying LD in individuals within different studies and populations result in the true causal variant being tagged by different SNPs in different people and studies. Several studies described here show altered expression of *OAS1* associated with these SNPs. Another possibility that has been proposed is that different SNPs within this locus differentially regulate the expression of genes within this cluster in different immune cells based on chromatin conformation (Schmiedel et al., 2020), although further work needs to be done to test the effects of different variants in other immune cell types including microglia and other tissue-specific macrophages, such as alveolar macrophages.

Lowering the expression of *OAS1* using siRNA to mimic the effects of an eQTL demonstrated that *OAS1* is required to dampen the expression and release of pro-inflammatory marker TNF-α when the levels of IFN-γ increase. Thus, individuals with eQTL variants that result in lower levels of *OAS1* expression may show more severe disease phenotypes associated with both AD and COVID-19, as a result of increased TNF-α and other pro-inflammatory cytokines. TNF-α and IFN-γ have been shown to have a synergistic effect on cell death (Karki et al., 2020), and so could damage nearby neurons and alveolar cells. TNF-α with C1Q complement (which also shows elevated expression in our h-iPSC-Mg system in response to *OAS1* knockdown), has been shown to induce the activation of reactive astrocytes, which contribute to synaptic damage, phagocytosis, and death of neurons and oligodendrocytes (Liddelow et al., 2017). Previously, we and others have shown that pro-inflammatory signals also suppress the expression of *Trem2* via TLR4 receptors (Liu et al., 2020; Owens et al., 2017; Zheng et al., 2016; Zhou et al., 2019). Our data is consistent with the suppression of *TREM2* expression as a response to IFN-γ and TNF-α in h-iPSC-Mg. Given that TREM2 is likely to have a protective role in AD and slow disease progression, elevated pro-inflammatory signals could increase the risk and progression of AD by suppressing this protective signal. TNF-α also activates PKR/ EIF2αK2, which is downstream of interferon signalling via TLR4, and consequently the protein activator of the IFN-inducible protein kinase (PRKRA) leading to apoptosis (Moradi Majd et al., 2020; Sadler and Williams, 2008). Altogether, chronic elevation of IFN signalling and subsequent TNF-α release could contribute to the development of dementia.

Pseudotime and network analysis of microglia isolated from *APP^NL-G-F^* and wild-type C57BL/6J mice across a variety of ages identified that *Oas1a* was upregulated alongside a co-expression network associated with interferon and pro-inflammatory responses expressed by a distinct sub-population of microglia transitioning to the interferon-related IRM state. This network contained *Oas1a* (the mouse orthologue of *OAS1* with 68% identity at the protein level), other *Oas* family members, *Mx1*, *Stat1/2, Ifit3, Ifitm3* and *Usp18* as hub genes. The expression of this network of genes was increased with age in wild-type mice. This finding suggests that the upregulation of interferon-responsive genes with age might mitigate the age-related damage by limiting pro-inflammatory signalling. Any dysfunction due to genetic variants in interferon signalling could result in persistent pro-inflammatory signalling and/or suppression of protective genes such as *TREM2* which could lead to dementia or severe COVID-19. Age is one of the strongest risk factors for severe COVID-19 responses and AD (Guerreiro and Bras, 2015; Ou et al., 2020), and so understanding the function of this genetic network will be important to attenuate the age-dependent contribution to these diseases.

The interferon-responsive genes seen in aged mice, and mice with amyloid plaques, were also expressed in a specific sub-population of microglia that are present in brains from people with AD (Olah et al. 2020). The genetic network we identified in human brains with AD containing *OAS1* and other interferon-responsive genes, identified *IFITM3*, *CD163*, and *ISG15* as hub genes. CD163 is a marker of a subset of border-associated macrophages (BAMs), that are present at the borders of the brain, including the meninges and choroid plexus, and is thought to represent an anti-inflammatory state in these cells, indicating the resolution of inflammation (Dvir-Szternfeld et al., 2021; Pey et al., 2014; De Schepper et al., 2020; Zhang et al., 2011). Indeed, emerging evidence suggests that SARS-CoV-2 is able to infect the brain vasculature, meninges, and choroid plexus (Yang et al., 2020), which would result in activation of BAMs and alter the cytokine environment around the brain leading to BBB breakdown, infiltration of immune cells into the brain, activation of astrocytes and phagocytosis of neurons. Coupling these findings with the high incidence of neurological problems in people showing a severe COVID-19 response (Mao et al., 2020; Paterson et al., 2020; Romero-Sánchez et al., 2020; Woo et al., 2020), emphasises the urgency with which we need to understand the mechanisms underlying the changes to cytokine signalling regulated by interferon signalling, particularly to anticipate the long-term neurological consequences of COVID-19 (Wang et al., 2020a). In addition to COVID-19, a SNP within *OAS1* was also associated with type 1 diabetes and so *OAS1* may integrate multiple risk factors that contribute to critical outcomes with COVID-19 (Tessier et al., 2006).

Surveying the genetic network we have identified containing *OAS1*, there are a number of secreted factors that may be useful as a readout of the state of these specialised innate immune cells in blood or CSF. The secreted genes include: *CXCL14, EGFL7, F13A1, ISG15, SCRG1, TGFBI* and *TNFSF13B*. The identification of *OAS1* and *OAS3* as genes that coordinate the response to amyloid-β plaques and COVID-19 infection poses the use of phosphodiesterase 12 (PDE12) inhibitors to increase the activity of OAS enzymes for these diseases, since PDE12 degrades the oligoadenylate activator of the OAS/RNase L system (Wood et al., 2015). PDE12 inhibitor compounds or *PDE12* knockout increase the anti-viral activity of cells. Furthermore, administration of interferons with correct timing may also help to dampen the pro-inflammatory response of innate immune cells, as in the treatment of multiple sclerosis (Davoudi-Monfared et al., 2020; Kingwell et al., 2019; Lee and Shin, 2020; Wang et al., 2020b).

In conclusion, our data show that OAS1 is required to limit the pro-inflammatory response of myeloid cells when stimulated with IFN-γ. We also identify a SNP within *OAS1* associated with AD in the same locus that predisposes to critical illness with COVID-19. This SNP acts as an eQTL, and is common in the population, and so may contribute to the high incidence of AD or critical illness with COVID-19 in the population. Further investigation of the function of OAS1 in innate immune cells and the genetic network engaged by *OAS1* will provide better molecular targets to track disease progression and treat AD, as well as COVID-19 and potentially its long-term sequelae.

## Acknowledgements

NM was supported by Alzheimer Nederland and the Erasmus+ Traineeship program. TMP was supported by funding to JMP and JH from the Innovative Medicines Initiative 2 Joint Undertaking under grant agreement 115976. This Joint Undertaking receives support from the European Union’s Horizon 2020 research and innovation program and the European Federation of Pharmaceutical Industries and Associations (EFPIA). RHR was supported through the award of a Leonard Wolfson Doctoral Training Fellowship in Neurodegeneration. JAB is supported through the Science and Technology Agency, Séneca Foundation, CARM, Spain (research project 00007/COVI/20). The University of Nottingham Group is funded by ARUK and hosts the ARUK Consortium DNA Bank, with the members given in the Appendix below. JH is supported by the Dolby Foundation, and by the National Institute for Health Research University College London Hospitals Biomedical Research Centre. DAS also received funding from the Alzheimer’s Research UK (ARUK) pump priming scheme via the UCL network. This work was funded by the UK DRI, which receives its funding from the DRI Ltd, funded by the UK Medical Research Council, Alzheimer’s Society and ARUK.

## Appendix

The University of Nottingham Group is funded by ARUK and hosts the ARUK Consortium DNA Bank, with the members: Tulsi Patel^1^, David M. Mann^2^, Peter Passmore^3^, David Craig^3^, Janet Johnston^3^, Bernadette McGuinness^3^, Stephen Todd^3^, Reinhard Heun^4^, Heike Kölsch^5^, Patrick G. Kehoe^6^, Emma R.L.C. Vardy^7^, Nigel M. Hooper^2^, Stuart Pickering-Brown^2^, Julie Snowden^8^, Anna Richardson^8^, Matt Jones^8^, David Neary^8^, Jenny Harris^8^, A. David Smith^9^, Gordon Wilcock^9^, Donald Warden^9^ and Clive Holmes^10^

^1^Schools of Life Sciences and Medicine, University of Nottingham, Nottingham NG7 2UH, UK, ^2^Institute of Brain, Behaviour and Mental Health, Faculty of Medical and Human Sciences, University of Manchester, Manchester M13 9PT, UK, ^3^Centre for Public Health, School of Medicine, Queen’s University Belfast, BT9 7BL, UK, ^4^Royal Derby Hospital, Derby DE22 3WQ, UK ^5^Department of Psychiatry, University of Bonn, Bonn 53105, Germany, ^6^School of Clinical Sciences, John James Laboratories, University of Bristol, Bristol BS16 1LE, UK, ^7^Salford Royal NHS Foundation Trust, ^8^Cerebral Function Unit, Greater Manchester Neurosciences Centre, Salford Royal Hospital, Stott Lane, Salford M6 8HD, UK, ^9^University of Oxford (OPTIMA), Oxford OX3 9DU, UK ^10^Clinical and Experimental Science, University of Southampton, Southampton SO17 1BJ, UK.

