## Supplementary data for "Genetic variability associated with *OAS1* expression in myeloid cells increases the risk of Alzheimer’s disease and severe COVID-19 outcomes"

### Supplementary Figures

**FIGURE S1. *OAS1* knockdown of h-iPSC-Mg in basal conditions showed a trend of increase in the expression of pro-inflammatory marker *TNF*.** (A) *OAS1* knockdown was mediated by using two siRNA sequences (1 and 2) targeting *OAS1*. One-way ANOVA showed a significant effect of siRNA treatment (p<0.01), indicated as a red line. (B-C) Pro-inflammatory marker *TNF* showed a trend of increase with *OAS1* knockdown; one-way ANOVA showed a trend for siRNA treatment (p=0.056), whereas for *IL1B* one-way ANOVA showed no significant effect of siRNA treatment (p>0.05). (D) The expression of *TGFB1* as an anti-inflammatory signalling marker, exhibited no change with *OAS1* knockdown. (E) The expression of *TREM2* was reduced in cells with *OAS1* knockdown mediated with siRNA-2. One-way ANOVA showed a significant effect of siRNA treatment (p=0.035), indicated as a horizontal red line. (F) A marker of microglial phagocytic activity, *CD68*, showed no significant change in the expression. (G-I) The markers of homeostatic microglia *C1QA*, *P2RY12* and *ITGAM* showed no detectable effect of *OAS1* knockdown. (J) Interferon-responsive *IFITM3* expression displayed a significant effect of siRNA treatment by one-way ANOVA (p=0.029), indicated as a horizontal red line. A significant increase in *IFITM3* was seen in cells with *OAS1* knockdown mediated with siRNA-2. (K-L) The expression of genes from the mouse or human interferon-response network, *STAT1* and *CD163*, showed no significant change with *OAS1* knockdown. Gene expression levels were normalised to the geometric mean of *GAPDH*, *HPRT1* and *RPS18*, then calculated as fold change relative to the non-targeting siRNA control (NC) in each individual culture preparation. N=5 independent plates. Data shown as mean ± SEM. One-way ANOVA; main effect of siRNA treatment indicated by horizontal red lines. When a significant main effect of siRNA treatment was seen, Dunnett's multiple comparisons tests were then performed to test for pairwise significance of siRNA-1 and -2 groups compared to the NC; * p<0.05, ** p<0.01, *** p<0.001, **** p<0.0001.

**FIGURE S2. The effects of IFN-γ treatment in h-iPSC-Mg with *OAS1* knockdown testing a series of markers.** (A) The expression of *IL1B* was not affected by IFN-γ treatment in *OAS1* knockdown iPSC-Mg. (B) The *TGFB1* anti-inflammatory marker showed significant upregulation in response to IFN-γ treatment regardless of *OAS1* knockdown. Two-way ANOVA with significant main effect of only IFN-γ treatment ($p<0.001$) indicated as a horizontal red line, no effect of siRNA treatment ($p>0.05$), and no significant interaction ($p>0.05$). (C) The downregulation of *TREM2* was observed with IFN-γ treated cells with no regard to *OAS1* knockdown. Two-way ANOVA with significant main effect of only IFN-γ treatment ($p<0.01$) indicated as a horizontal red line, no effect of siRNA treatment ($p>0.05$), and no significant interaction ($p>0.05$). (D) The slight upregulation in *CD68* expression was detected in IFN-γ treated cells independently of *OAS1* knockdown. Two-way ANOVA with significant main effect of only IFN-γ treatment ($p<0.01$) indicated as a horizontal red line, no effect of siRNA treatment ($p>0.05$), and no significant interaction ($p>0.05$). (E) The significant upregulation of *P2RY12* was observed in response to IFN-γ treatment regardless of *OAS1* knockdown. Two-way ANOVA with significant main effect of only IFN-γ treatment ($p<0.01$) indicated as a horizontal red line, no effect of siRNA treatment ($p>0.05$), and no significant interaction ($p>0.05$). (F) No significant effect of IFN-γ treatment was detected for *ITGAM* expression. Two-way ANOVA with significant main effect of only siRNA treatment ($p<0.01$) indicated as a vertical red line, no effect of IFN-γ treatment ($p>0.05$), and a significant interaction ($p=0.027$). (G) The expression of *IFITM3* was upregulated in response to IFN-γ treatment independently from *OAS1* knockdown. Two-way ANOVA with significant main effect of only IFN-γ treatment ($p<0.01$) indicated as a horizontal red line, no effect of siRNA treatment ($p>0.05$), and no significant interaction ($p>0.05$). (H) The upregulation of *STAT1* expression in cells treated with IFN-γ was observed with no regard to *OAS1* knockdown. Two-way ANOVA with significant main effect of only IFN-γ treatment ($p=0.035$) indicated as a horizontal red line, no effect of siRNA treatment ($p>0.05$), and no significant interaction ($p>0.05$). (I) The expression of *CD163* was not affected by IFN-γ treatment in cells with *OAS1* knockdown. Gene expression levels were normalised to the geometric mean of *GAPDH, HPRT1* and *RPS18,* then calculated as fold change relative to the non-targeting siRNA control (vehicle control) without IFN-γ treatment in each individual culture preparation. N=5 independent plates. Data shown as mean ± SEM. Two-way ANOVA; significant main effects of IFN-γ treatment and *OAS1* knockdown indicated by red horizontal and vertical lines respectively. When a significant interaction was seen between IFN-γ treatment and siRNA treatment, Tukey's multiple comparisons tests were then performed to test for pairwise significance of all possible groups; * $p<0.05$, ** $p<0.01$, *** $p<0.001$, **** $p<0.0001$.

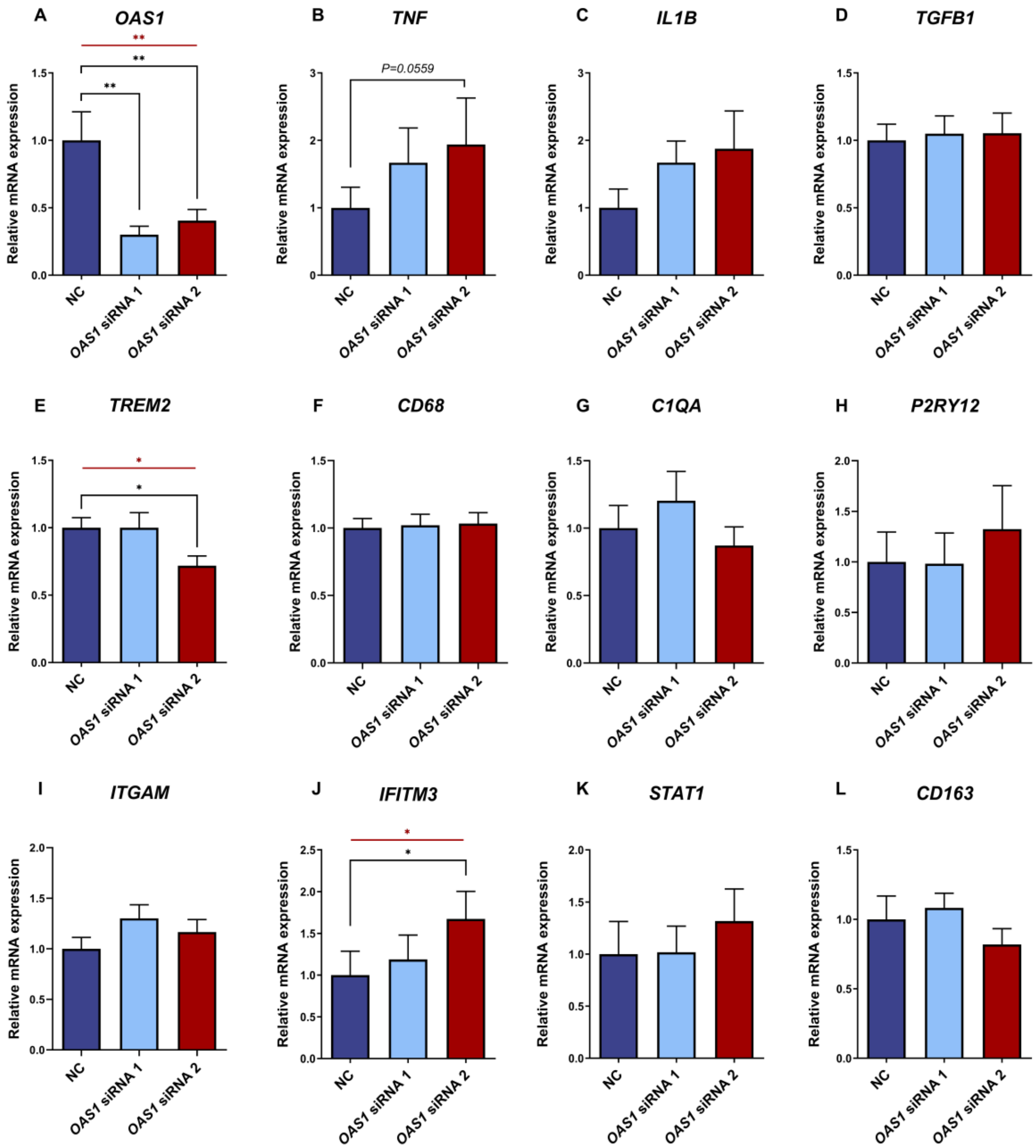
A
OAS1
B
TNF
P=0.0559
C
IL1B
D
TGFB1
E
TREM2
F
CD68
G
C1QA
H
P2RY12
I
ITGAM
J
IFITM3
K
STAT1
L
CD163
Relative mRNA expression
NC
OAS1 siRNA 1
OAS1 siRNA 2

**Fig. S1**

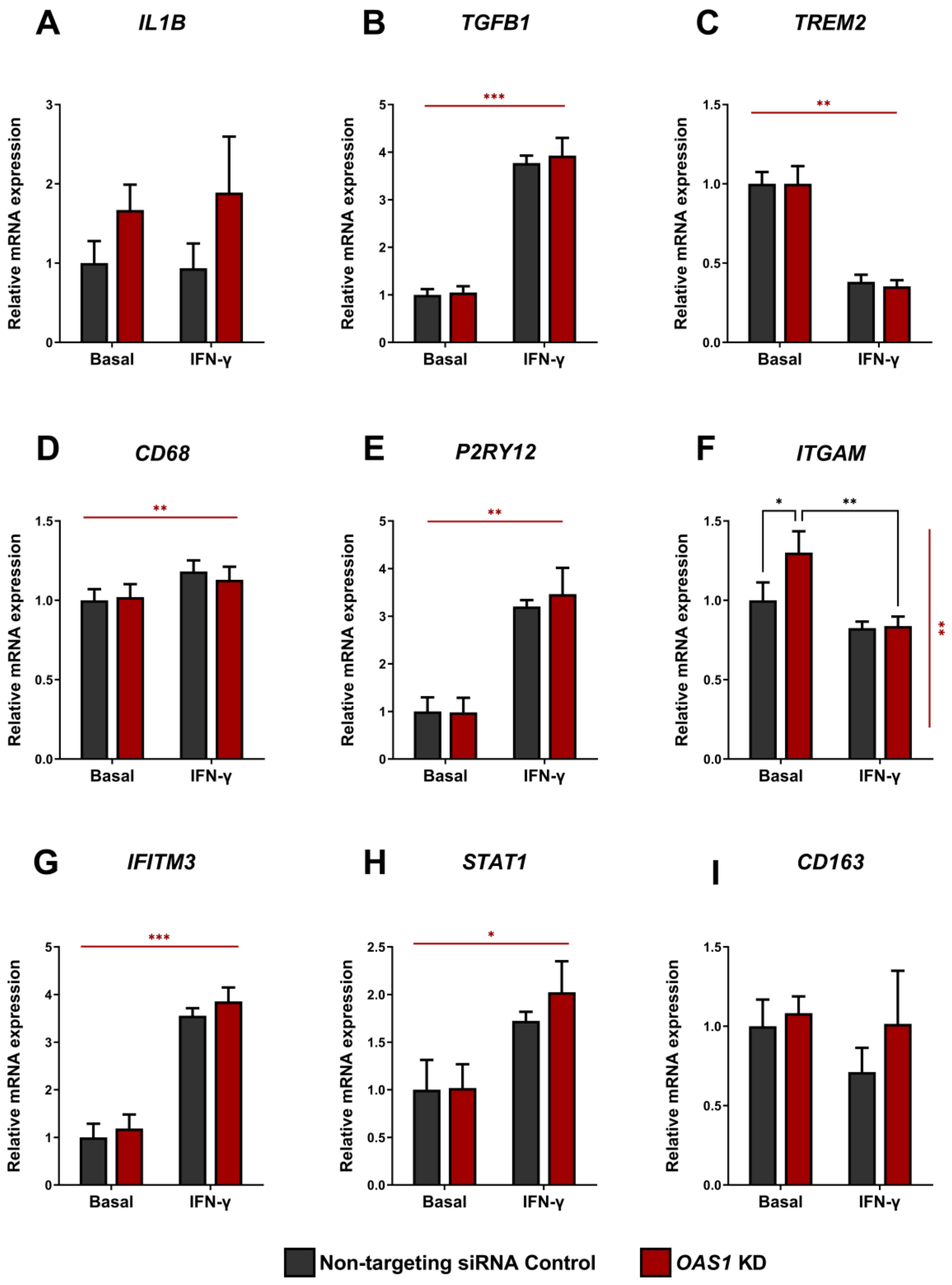
A
IL1B
B
TGFB1
C
TREM2
D
CD68
E
P2RY12
F
ITGAM
G
IFITM3
H
STAT1
I
CD163
Relative mRNA expression
Basal
IFN-γ
Non-targeting siRNA Control
OAS1 KD

Fig. S2

**Table S1 Primers sequences used for RT-qPCR**

| Target Gene | Forward Primer (5'->3') | Reverse Primer (5'->3') | Source |
|---|---|---|---|
| *C1QA* | GAAGAAAGGGGAGGCAGGAAGAC | TGCCCTTGGTGCCTTTAATTCC | Sigma Merck |
| *CD163* | TCCTGTAACTGCTCTAGGTGC | TCTCTACTCTCCCAGCACAGC | Sigma Merck |
| *CD68* | GACCTCCAGCAGAAGGTTGTC | GAGGTGGACAGCTGGTGAAAG | Sigma Merck |
| *GAPDH* | GCCAAAAGGGTCATCATCTCTG | CAGTCTTCTGGGTGGCAGTG | Sigma Merck |
| *HPRT1* | CTTTGCTTTCCTTGGTCAGGC | TATATCCAACACTTCGTGGGGTC | Sigma Merck |
| *IFITM3* | CATGTCGTCTGGTCCCTGTTC | ATCCATAGGCCTGGAAGATCAGC | Sigma Merck |
| *IL1B* | TGAAGCTGATGGCCCTAAACAG | AAGGTGCTCAGGTCATTCTCC | Sigma Merck |
| *ITGAM* | TCCAACGCTAATGTCAAGGGC | GGTCTGCTCGTAGTAATGGGGG | Sigma Merck |
| *OAS1* | AAGCCTGTCAAAGAGAGAGAGC | GGTTAGGTTTATAGCCGCCAG | Sigma Merck |
| *P2RY12* | ACCCTCCAGAATCAACAGTTATC | GTGTAGAGCAGTGGGAAGAGG | Sigma Merck |
| *RPS18* | GATGGGCGGCGGAAAATAGC | GGTCAATGTCTGCTTTCCTCAAC | Sigma Merck |
| *STAT1* | AGCTGTCTGAAGGAAGAAAGG | GTTCTGCAAGGTTTTGCATTTGAAG | Sigma Merck |
| *TGFB1* | ACAAGTTCAAGCAGAGTACACAC | CATCAAAAGATAACCACTCTGGCG | Sigma Merck |
| *TNF* | TCTCGAACCCCGAGTGACAAG | CTGGTTATCTCTCAGCTCCACG | Sigma Merck |
| *TREM2* | GGAGTCTGAGAGCTTCGAGGATG | TTCACTGGGTGGATGTGTCCC | Sigma Merck |

Table S2 Microglial interferon response module in APP-KI versus WT mice

| | ensgene | name | module | mm |
|---|---|---|---|---|
| 390 | Ifit3 | Ifit3 | magenta | 0.806958 |
| 188 | Ifitm3 | Ifitm3 | magenta | 0.740379 |
| 147 | Usp18 | Usp18 | magenta | 0.730901 |
| 32 | Ifi211 | Ifi211 | magenta | 0.717632 |
| 31 | Ifi204 | Ifi204 | magenta | 0.700952 |
| 189 | Irf7 | Irf7 | magenta | 0.700632 |
| 389 | Ifit2 | Ifit2 | magenta | 0.694785 |
| 25 | Ifi206 | Ifi206 | magenta | 0.683875 |
| 118 | Oasl2 | Oasl2 | magenta | 0.677706 |
| 391 | Ifit3b | Ifit3b | magenta | 0.676638 |
| 249 | Tgtp2 | Tgtp2 | magenta | 0.66573 |
| 29 | Ifi208 | Ifi208 | magenta | 0.66216 |
| 392 | Ifit1 | Ifit1 | magenta | 0.659513 |
| 376 | Iigp1 | Iigp1 | magenta | 0.638397 |
| 124 | Oas1a | Oas1a | magenta | 0.630879 |
| 30 | Ifi207 | Ifi207 | magenta | 0.630743 |
| 280 | Rsad2 | Rsad2 | magenta | 0.622057 |
| 332 | Rtp4 | Rtp4 | magenta | 0.598434 |
| 119 | Oasl1 | Oasl1 | magenta | 0.595185 |
| 120 | Oas2 | Oas2 | magenta | 0.594603 |
| 387 | RP24-84E1 | RP24-84E1 | magenta | 0.590712 |
| 260 | Ccl12 | Ccl12 | magenta | 0.589702 |
| 27 | Ifi213 | Ifi213 | magenta | 0.583964 |
| 55 | Zbp1 | Zbp1 | magenta | 0.571114 |
| 310 | Phf11b | Phf11b | magenta | 0.564387 |
| 261 | Slfn5 | Slfn5 | magenta | 0.558635 |
| 342 | Mx1 | Mx1 | magenta | 0.549915 |
| 250 | Ifi47 | Ifi47 | magenta | 0.539879 |
| 28 | Ifi209 | Ifi209 | magenta | 0.511232 |
| 372 | Gm4951 | Gm4951 | magenta | 0.498206 |
| 123 | Oas1g | Oas1g | magenta | 0.492276 |
| 106 | Cxcl10 | Cxcl10 | magenta | 0.489592 |
| 281 | Cmpk2 | Cmpk2 | magenta | 0.487167 |
| 311 | Phf11d | Phf11d | magenta | 0.475521 |
| 4 | Stat1 | Stat1 | magenta | 0.461982 |
| 121 | Oas3 | Oas3 | magenta | 0.449652 |
| 112 | Gbp6 | Gbp6 | magenta | 0.442922 |
| 10 | Sp100 | Sp100 | magenta | 0.441394 |
| 343 | Mx2 | Mx2 | magenta | 0.433119 |
| 264 | Slfn2 | Slfn2 | magenta | 0.41988 |
| 207 | Nlrc5 | Nlrc5 | magenta | 0.419414 |
| 384 | Ms4a4c | Ms4a4c | magenta | 0.416574 |
| 239 | Stat2 | Stat2 | magenta | 0.405266 |
| 276 | Rnf213 | Rnf213 | magenta | 0.404272 |

| | | | | |
|---|---|---|---|---|
| 309 | Phf11a | Phf11a | magenta | 0.400679 |
| 201 | Bst2 | Bst2 | magenta | 0.399164 |
| 259 | Ccl2 | Ccl2 | magenta | 0.391116 |
| 288 | Ifi27l2a | Ifi27l2a | magenta | 0.387914 |
| 254 | Gm12250 | Gm12250 | magenta | 0.367725 |
| 222 | Pml | Pml | magenta | 0.349504 |
| 96 | Fgl2 | Fgl2 | magenta | 0.347956 |
| 257 | Xaf1 | Xaf1 | magenta | 0.343483 |
| 166 | Isg20 | Isg20 | magenta | 0.343417 |
| 245 | Irgm1 | Irgm1 | magenta | 0.321676 |
| 17 | Tor3a | Tor3a | magenta | 0.321204 |
| 335 | Parp14 | Parp14 | magenta | 0.320177 |
| 270 | Dhx58 | Dhx58 | magenta | 0.316204 |
| 8 | Gm7609 | Gm7609 | magenta | 0.315451 |
| 36 | Ifih1 | Ifih1 | magenta | 0.312142 |
| 57 | Helz2 | Helz2 | magenta | 0.30926 |
| 319 | Ly6e | Ly6e | magenta | 0.30834 |
| 255 | Igtp | Igtp | magenta | 0.306926 |
| 375 | F830016B( | F830016B( | magenta | 0.305152 |
| 75 | Ddx58 | Ddx58 | magenta | 0.305019 |
| 175 | Trim30a | Trim30a | magenta | 0.302953 |
| 111 | Gbp4 | Gbp4 | magenta | 0.286582 |
| 9 | Gm7592 | Gm7592 | magenta | 0.283771 |
| 374 | Gm4841 | Gm4841 | magenta | 0.281893 |
| 70 | Gbp2 | Gbp2 | magenta | 0.276488 |
| 321 | Ly6a | Ly6a | magenta | 0.269294 |
| 248 | Tgtp1 | Tgtp1 | magenta | 0.268689 |
| 308 | Gm6904 | Gm6904 | magenta | 0.268172 |
| 303 | BC147527 | BC147527 | magenta | 0.265169 |
| 330 | Socs1 | Socs1 | magenta | 0.263766 |
| 366 | Eif2ak2 | Eif2ak2 | magenta | 0.260287 |
| 61 | Fcgr1 | Fcgr1 | magenta | 0.259932 |
| 262 | Slfn9 | Slfn9 | magenta | 0.257266 |
| 197 | Gm45418 | Gm45418 | magenta | 0.257007 |
| 357 | H2-T23 | H2-T23 | magenta | 0.255015 |
| 263 | Slfn8 | Slfn8 | magenta | 0.252366 |
| 87 | Sdc3 | Sdc3 | magenta | 0.250575 |
| 134 | Gm20559 | Gm20559 | magenta | 0.24833 |
| 177 | Trim30d | Trim30d | magenta | 0.246359 |
| 37 | Ube2l6 | Ube2l6 | magenta | 0.245259 |
| 381 | Batf2 | Batf2 | magenta | 0.244184 |
| 265 | Ccl5 | Ccl5 | magenta | 0.243043 |
| 71 | Ifi44 | Ifi44 | magenta | 0.242205 |
| 135 | Samd9l | Samd9l | magenta | 0.24063 |
| 122 | Oas1b | Oas1b | magenta | 0.240374 |
| 179 | Gm1966 | Gm1966 | magenta | 0.238335 |
| 354 | H2-Q6 | H2-Q6 | magenta | 0.237829 |

| | | | | |
|---|---|---|---|---|
| 246 | Gm5431 | Gm5431 | magenta | 0.236153 |
| 362 | B430306N | B430306N | magenta | 0.232101 |
| 337 | Parp9 | Parp9 | magenta | 0.231344 |
| 178 | Gm8995 | Gm8995 | magenta | 0.230874 |
| 7 | Gm15433 | Gm15433 | magenta | 0.229742 |
| 60 | Adar | Adar | magenta | 0.227013 |
| 355 | H2-Q7 | H2-Q7 | magenta | 0.224137 |
| 58 | Tnfsf10 | Tnfsf10 | magenta | 0.222703 |
| 52 | Znfx1 | Znfx1 | magenta | 0.222469 |
| 272 | Ifi35 | Ifi35 | magenta | 0.221018 |
| 138 | Parp12 | Parp12 | magenta | 0.214454 |
| 141 | Herc6 | Herc6 | magenta | 0.213976 |
| 174 | Trim30c | Trim30c | magenta | 0.213257 |
| 251 | Irf1 | Irf1 | magenta | 0.207551 |
| 256 | Scimp | Scimp | magenta | 0.206561 |
| 282 | Nampt | Nampt | magenta | 0.206025 |
| 373 | Gm5970 | Gm5970 | magenta | 0.203247 |
| 125 | Trafd1 | Trafd1 | magenta | 0.200854 |
| 69 | Gbp3 | Gbp3 | magenta | 0.199879 |
| 26 | Ifi214 | Ifi214 | magenta | 0.198581 |
| 352 | C4b | C4b | magenta | 0.19473 |
| 312 | Phf11c | Phf11c | magenta | 0.19302 |
| 88 | Clic4 | Clic4 | magenta | 0.189415 |
| 190 | Slc25a22 | Slc25a22 | magenta | 0.187457 |
| 356 | H2-T24 | H2-T24 | magenta | 0.186753 |
| 258 | Lgals9 | Lgals9 | magenta | 0.186528 |
| 51 | Samhd1 | Samhd1 | magenta | 0.186452 |
| 173 | Trim30b | Trim30b | magenta | 0.185478 |
| 320 | Ly6i | Ly6i | magenta | 0.18505 |
| 21 | Fcgr4 | Fcgr4 | magenta | 0.185029 |
| 347 | Daxx | Daxx | magenta | 0.183356 |
| 47 | Siglec1 | Siglec1 | magenta | 0.181453 |
| 230 | Zufsp | Zufsp | magenta | 0.181311 |
| 110 | Gbp9 | Gbp9 | magenta | 0.179655 |
| 132 | Tpst1 | Tpst1 | magenta | 0.179574 |
| 277 | Gm44935 | Gm44935 | magenta | 0.177791 |
| 104 | Dck | Dck | magenta | 0.177606 |
| 323 | Parp10 | Parp10 | magenta | 0.174988 |
| 305 | Psme1 | Psme1 | magenta | 0.174682 |
| 198 | Ddx60 | Ddx60 | magenta | 0.173797 |
| 266 | Trim25 | Trim25 | magenta | 0.173558 |
| 38 | Prr5l | Prr5l | magenta | 0.172422 |
| 64 | Vcam1 | Vcam1 | magenta | 0.172094 |
| 67 | Gbp5 | Gbp5 | magenta | 0.171865 |
| 56 | Ogfr | Ogfr | magenta | 0.169928 |
| 137 | Zc3hav1 | Zc3hav1 | magenta | 0.166227 |
| 315 | Epsti1 | Epsti1 | magenta | 0.163222 |

| | | | | |
|---|---|---|---|---|
| 328 | Gpr84 | Gpr84 | magenta | 0.151026 |
| 92 | Mthfr | Mthfr | magenta | 0.150192 |
| 350 | Tap1 | Tap1 | magenta | 0.150064 |
| 361 | Treml2 | Treml2 | magenta | 0.144717 |
| 24 | Slamf8 | Slamf8 | magenta | 0.143125 |
| 268 | Hap1 | Hap1 | magenta | 0.141439 |
| 383 | Ccdc86 | Ccdc86 | magenta | 0.138123 |
| 206 | 9330175E1 | 9330175E1 | magenta | 0.137243 |
| 34 | Nmi | Nmi | magenta | 0.137103 |
| 395 | Pik3ap1 | Pik3ap1 | magenta | 0.135975 |
| 162 | 1600014C1 | 1600014C1 | magenta | 0.134428 |
| 154 | Etnk1 | Etnk1 | magenta | 0.132595 |
| 204 | Gm17435 | Gm17435 | magenta | 0.132584 |
| 15 | Tor1aip1 | Tor1aip1 | magenta | 0.128592 |
| 68 | Gbp7 | Gbp7 | magenta | 0.126992 |
| 225 | Shisa5 | Shisa5 | magenta | 0.126515 |
| 238 | Timeless | Timeless | magenta | 0.122181 |
| 349 | Psmb9 | Psmb9 | magenta | 0.120891 |
| 223 | Gm2065 | Gm2065 | magenta | 0.120292 |
| 306 | Irf9 | Irf9 | magenta | 0.120127 |
| 405 | Chic1 | Chic1 | magenta | 0.119371 |
| 12 | Mybph | Mybph | magenta | 0.119347 |
| 195 | Tlr3 | Tlr3 | magenta | 0.117972 |
| 359 | Gm20429 | Gm20429 | magenta | 0.1178 |
| 327 | Apobec3 | Apobec3 | magenta | 0.11664 |
| 53 | Zfas1 | Zfas1 | magenta | 0.113614 |
| 131 | Tctn2 | Tctn2 | magenta | 0.11344 |
| 172 | Trim12c | Trim12c | magenta | 0.111582 |
| 336 | Dtx3l | Dtx3l | magenta | 0.110777 |
| 126 | Naa25 | Naa25 | magenta | 0.109239 |
| 313 | Setdb2 | Setdb2 | magenta | 0.109079 |
| 252 | Gm12216 | Gm12216 | magenta | 0.108015 |
| 151 | 2310001H | 2310001H | magenta | 0.107672 |
| 196 | Sap30 | Sap30 | magenta | 0.106336 |
| 348 | Tapbp | Tapbp | magenta | 0.106143 |
| 2 | Arid5a | Arid5a | magenta | 0.10599 |
| 353 | Tnf | Tnf | magenta | 0.10596 |
| 283 | Gdap10 | Gdap10 | magenta | 0.105887 |
| 139 | Creb5 | Creb5 | magenta | 0.105038 |
| 107 | Cxcl13 | Cxcl13 | magenta | 0.104403 |
| 247 | Gm12185 | Gm12185 | magenta | 0.103479 |
| 65 | Sass6 | Sass6 | magenta | 0.103293 |
| 40 | Thbs1 | Thbs1 | magenta | 0.099417 |
| 95 | AW01173 | AW01173 | magenta | 0.09802 |
| 221 | Pstpip1 | Pstpip1 | magenta | 0.097148 |
| 214 | Casp4 | Casp4 | magenta | 0.095903 |
| 217 | Icam1 | Icam1 | magenta | 0.09584 |

| | | | | |
|---|---|---|---|---|
| 278 | Baiap2 | Baiap2 | magenta | 0.09527 |
| 101 | Lap3 | Lap3 | magenta | 0.093738 |
| 340 | Dopey2 | Dopey2 | magenta | 0.090134 |
| 244 | Pttg1 | Pttg1 | magenta | 0.088223 |
| 140 | Nod1 | Nod1 | magenta | 0.086891 |
| 229 | Ncoa7 | Ncoa7 | magenta | 0.086626 |
| 325 | Tmem184l | Tmem184l | magenta | 0.08593 |
| 169 | Nup98 | Nup98 | magenta | 0.084644 |
| 358 | H2-T22 | H2-T22 | magenta | 0.084252 |
| 44 | Sppl2a | Sppl2a | magenta | 0.084221 |
| 148 | Apobec1 | Apobec1 | magenta | 0.084071 |
| 130 | Rilpl1 | Rilpl1 | magenta | 0.084049 |
| 304 | Rnase6 | Rnase6 | magenta | 0.0836 |
| 54 | Rnf114 | Rnf114 | magenta | 0.083003 |
| 269 | Cnp | Cnp | magenta | 0.080799 |
| 108 | Rasgef1b | Rasgef1b | magenta | 0.080034 |
| 385 | Ms4a6c | Ms4a6c | magenta | 0.078373 |
| 267 | Arl5c | Arl5c | magenta | 0.075856 |
| 199 | Comp | Comp | magenta | 0.075626 |
| 402 | 5430427O | 5430427O | magenta | 0.074822 |
| 18 | 4930523C( | 4930523C( | magenta | 0.074025 |
| 202 | Tmem221 | Tmem221 | magenta | 0.073559 |
| 393 | Pcgf5 | Pcgf5 | magenta | 0.07279 |
| 284 | Nfkbia | Nfkbia | magenta | 0.072308 |
| 301 | Cd180 | Cd180 | magenta | 0.072107 |
| 294 | Fbxw17 | Fbxw17 | magenta | 0.0714 |
| 213 | Casp1 | Casp1 | magenta | 0.070905 |
| 22 | Nectin4 | Nectin4 | magenta | 0.0706 |
| 48 | Xrn2 | Xrn2 | magenta | 0.070397 |
| 360 | Enpp4 | Enpp4 | magenta | 0.070147 |
| 168 | Acer3 | Acer3 | magenta | 0.069953 |
| 159 | Sertad3 | Sertad3 | magenta | 0.069659 |
| 386 | BE692007 | BE692007 | magenta | 0.069571 |
| 86 | Marcksl1 | Marcksl1 | magenta | 0.069494 |
| 155 | 4930479D | 4930479D | magenta | 0.069269 |
| 81 | Slc31a2 | Slc31a2 | magenta | 0.066967 |
| 334 | Gm17106 | Gm17106 | magenta | 0.066753 |
| 105 | Naaa | Naaa | magenta | 0.066234 |
| 184 | Il27 | Il27 | magenta | 0.066232 |
| 1 | Sgk3 | Sgk3 | magenta | 0.065558 |
| 329 | Nmral1 | Nmral1 | magenta | 0.065288 |
| 241 | Peli1 | Peli1 | magenta | 0.065204 |
| 369 | Map3k8 | Map3k8 | magenta | 0.06464 |
| 351 | Tap2 | Tap2 | magenta | 0.063122 |
| 215 | Naalad2 | Naalad2 | magenta | 0.062224 |
| 3 | Mitd1 | Mitd1 | magenta | 0.061834 |
| 82 | Slc31a1 | Slc31a1 | magenta | 0.061737 |

| | | | | |
|---|---|---|---|---|
| 133 | Rasa4 | Rasa4 | magenta | 0.061452 |
| 153 | St8sia1 | St8sia1 | magenta | 0.061096 |
| 43 | Spint1 | Spint1 | magenta | 0.060824 |
| 399 | Cfap43 | Cfap43 | magenta | 0.06044 |
| 326 | Gm16576 | Gm16576 | magenta | 0.060344 |
| 403 | Igbp1 | Igbp1 | magenta | 0.060045 |
| 404 | Nono | Nono | magenta | 0.059881 |
| 295 | Sema4d | Sema4d | magenta | 0.059756 |
| 33 | Wdr38 | Wdr38 | magenta | 0.059198 |
| 345 | 9530082P2 | 9530082P2 | magenta | 0.059074 |
| 103 | Tmem156 | Tmem156 | magenta | 0.059073 |
| 339 | Usp25 | Usp25 | magenta | 0.05763 |
| 116 | Ankle2 | Ankle2 | magenta | 0.057326 |
| 371 | Dcp2 | Dcp2 | magenta | 0.057163 |
| 271 | Kat2a | Kat2a | magenta | 0.056594 |
| 243 | Pnpt1 | Pnpt1 | magenta | 0.055832 |
| 79 | Tdrd7 | Tdrd7 | magenta | 0.055505 |
| 183 | Gm14388 | Gm14388 | magenta | 0.055285 |
| 150 | Clec2d | Clec2d | magenta | 0.054758 |
| 365 | Gm37639 | Gm37639 | magenta | 0.054395 |
| 167 | Gm26522 | Gm26522 | magenta | 0.053552 |
| 78 | Zbtb5 | Zbtb5 | magenta | 0.052998 |
| 233 | Plpp2 | Plpp2 | magenta | 0.052637 |
| 203 | Klf2 | Klf2 | magenta | 0.051575 |
| 14 | Rnpep | Rnpep | magenta | 0.051283 |
| 370 | Snhg4 | Snhg4 | magenta | 0.051096 |
| 171 | Trim21 | Trim21 | magenta | 0.050726 |
| 13 | Ppfia4 | Ppfia4 | magenta | 0.050399 |
| 219 | Sik3 | Sik3 | magenta | 0.050023 |
| 59 | Selenot | Selenot | magenta | 0.049773 |
| 192 | Cd81 | Cd81 | magenta | 0.049733 |
| 406 | G530011O | G530011O | magenta | 0.049721 |
| 157 | Bcl3 | Bcl3 | magenta | 0.049301 |
| 377 | Tcof1 | Tcof1 | magenta | 0.04915 |
| 324 | Triobp | Triobp | magenta | 0.049066 |
| 182 | Gm45191 | Gm45191 | magenta | 0.049001 |
| 84 | Prpf38a | Prpf38a | magenta | 0.048427 |
| 50 | E130215H | E130215H | magenta | 0.047914 |
| 149 | Parp11 | Parp11 | magenta | 0.047349 |
| 367 | Rmdn2 | Rmdn2 | magenta | 0.047269 |
| 49 | Apmap | Apmap | magenta | 0.047191 |
| 289 | Asb13 | Asb13 | magenta | 0.047172 |
| 80 | Ugcg | Ugcg | magenta | 0.046959 |
| 185 | Tmem219 | Tmem219 | magenta | 0.046327 |
| 220 | Zc3h12c | Zc3h12c | magenta | 0.04594 |
| 73 | Cpne3 | Cpne3 | magenta | 0.044862 |
| 165 | Gm45109 | Gm45109 | magenta | 0.044685 |

| | | | | |
|---|---|---|---|---|
| 90 | Gm13212 | Gm13212 | magenta | 0.044627 |
| 35 | Fmnl2 | Fmnl2 | magenta | 0.044564 |
| 94 | Morn1 | Morn1 | magenta | 0.044353 |
| 176 | Gm25405 | Gm25405 | magenta | 0.044117 |
| 142 | Rab43 | Rab43 | magenta | 0.044105 |
| 333 | Il1rap | Il1rap | magenta | 0.044051 |
| 382 | Pla2g16 | Pla2g16 | magenta | 0.043834 |
| 226 | Katna1 | Katna1 | magenta | 0.043423 |
| 136 | Akr1b3 | Akr1b3 | magenta | 0.043262 |
| 180 | Cyp2r1 | Cyp2r1 | magenta | 0.043136 |
| 298 | Papd7 | Papd7 | magenta | 0.042847 |
| 170 | Pgap2 | Pgap2 | magenta | 0.042824 |
| 237 | Slc16a7 | Slc16a7 | magenta | 0.042419 |
| 216 | A230050P | A230050P | magenta | 0.042018 |
| 145 | Itpr1 | Itpr1 | magenta | 0.04156 |
| 232 | Ybey | Ybey | magenta | 0.041471 |
| 286 | Susd6 | Susd6 | magenta | 0.041447 |
| 394 | Btaf1 | Btaf1 | magenta | 0.040605 |
| 363 | Mocs1 | Mocs1 | magenta | 0.039966 |
| 322 | Naprt | Naprt | magenta | 0.038898 |
| 63 | Mov10 | Mov10 | magenta | 0.038845 |
| 146 | Csgalnact2 | Csgalnact2 | magenta | 0.038723 |
| 302 | Ppwd1 | Ppwd1 | magenta | 0.038598 |
| 285 | Daam1 | Daam1 | magenta | 0.03817 |
| 77 | Tesk1 | Tesk1 | magenta | 0.038142 |
| 114 | 5430403G | 5430403G | magenta | 0.038086 |
| 99 | Afap1 | Afap1 | magenta | 0.037928 |
| 91 | Zfp933 | Zfp933 | magenta | 0.037787 |
| 210 | Mthfsd | Mthfsd | magenta | 0.037773 |
| 143 | Kbtbd8 | Kbtbd8 | magenta | 0.037521 |
| 235 | Fbxo7 | Fbxo7 | magenta | 0.037348 |
| 398 | Sfxn2 | Sfxn2 | magenta | 0.037299 |
| 300 | Mrps27 | Mrps27 | magenta | 0.037119 |
| 16 | 9430034N | 9430034N | magenta | 0.036952 |
| 115 | Gm15446 | Gm15446 | magenta | 0.036352 |
| 89 | Fhad1 | Fhad1 | magenta | 0.03635 |
| 39 | Cstf3 | Cstf3 | magenta | 0.036016 |
| 193 | Abhd13 | Abhd13 | magenta | 0.035633 |
| 218 | RP23-162F | RP23-162F | magenta | 0.035366 |
| 45 | Gm10766 | Gm10766 | magenta | 0.034956 |
| 187 | Armc5 | Armc5 | magenta | 0.034688 |
| 117 | Tfip11 | Tfip11 | magenta | 0.034582 |
| 102 | Tbc1d1 | Tbc1d1 | magenta | 0.034571 |
| 93 | Cep104 | Cep104 | magenta | 0.034564 |
| 380 | Snx15 | Snx15 | magenta | 0.034473 |
| 152 | Ybx3 | Ybx3 | magenta | 0.034305 |
| 240 | Osm | Osm | magenta | 0.034165 |

| | | | | |
|---|---|---|---|---|
| 378 | Poli | Poli | magenta | 0.033557 |
| 11 | Armc9 | Armc9 | magenta | 0.03343 |
| 401 | Fam122b | Fam122b | magenta | 0.033361 |
| 46 | A730017L: | A730017L: | magenta | 0.033141 |
| 364 | Epb41l3 | Epb41l3 | magenta | 0.033026 |
| 74 | 3110043O | 3110043O | magenta | 0.032773 |
| 314 | Ccdc122 | Ccdc122 | magenta | 0.032744 |
| 231 | Nt5dc1 | Nt5dc1 | magenta | 0.032719 |
| 346 | Taf11 | Taf11 | magenta | 0.032247 |
| 98 | Letm1 | Letm1 | magenta | 0.03178 |
| 163 | Fam71e1 | Fam71e1 | magenta | 0.031695 |
| 85 | Zfyve9 | Zfyve9 | magenta | 0.031654 |
| 100 | Tmem128 | Tmem128 | magenta | 0.031646 |
| 164 | Ppp1r15a | Ppp1r15a | magenta | 0.030957 |
| 109 | Klhl8 | Klhl8 | magenta | 0.030903 |
| 253 | Anxa6 | Anxa6 | magenta | 0.030337 |
| 113 | Mfsd7a | Mfsd7a | magenta | 0.03021 |
| 297 | Ptch1 | Ptch1 | magenta | 0.030178 |
| 129 | Tctn1 | Tctn1 | magenta | 0.030066 |
| 291 | Ripk1 | Ripk1 | magenta | 0.029912 |
| 42 | Dnajc17 | Dnajc17 | magenta | 0.02978 |
| 400 | Enox2 | Enox2 | magenta | 0.029779 |
| 5 | Inpp1 | Inpp1 | magenta | 0.029698 |
| 368 | Atl2 | Atl2 | magenta | 0.028835 |
| 242 | Pus10 | Pus10 | magenta | 0.027692 |
| 127 | Acad10 | Acad10 | magenta | 0.02748 |
| 331 | 3110001I2 | 3110001I2 | magenta | 0.027409 |
| 234 | RP24-406I | RP24-406I | magenta | 0.027346 |
| 212 | 2610044O | 2610044O | magenta | 0.027244 |
| 318 | Trappc9 | Trappc9 | magenta | 0.027104 |
| 274 | Socs3 | Socs3 | magenta | 0.027092 |
| 6 | Atic | Atic | magenta | 0.026613 |
| 23 | Slamf6 | Slamf6 | magenta | 0.026596 |
| 317 | Nudcd1 | Nudcd1 | magenta | 0.026567 |
| 160 | Nfkbid | Nfkbid | magenta | 0.026224 |
| 397 | Psd | Psd | magenta | 0.026213 |
| 338 | Arl13b | Arl13b | magenta | 0.026046 |
| 211 | Gas8 | Gas8 | magenta | 0.024947 |
| 388 | Asah2 | Asah2 | magenta | 0.024153 |
| 208 | Acd | Acd | magenta | 0.024131 |
| 83 | Cntln | Cntln | magenta | 0.024014 |
| 307 | Rcbtb1 | Rcbtb1 | magenta | 0.02376 |
| 66 | Npnt | Npnt | magenta | 0.022487 |
| 344 | Zfp946 | Zfp946 | magenta | 0.021618 |
| 205 | Rpgrip1l | Rpgrip1l | magenta | 0.021446 |
| 290 | Tdp2 | Tdp2 | magenta | 0.021337 |
| 62 | Rnf115 | Rnf115 | magenta | 0.021217 |

| | | | | |
|---|---|---|---|---|
| 293 | Mcur1 | Mcur1 | magenta | 0.021192 |
| 194 | Atp11a | Atp11a | magenta | 0.020418 |
| 299 | Ttc37 | Ttc37 | magenta | 0.020222 |
| 161 | Hpn | Hpn | magenta | 0.020152 |
| 224 | Nicn1 | Nicn1 | magenta | 0.020114 |
| 128 | Brap | Brap | magenta | 0.020017 |
| 209 | Wdr59 | Wdr59 | magenta | 0.019826 |
| 186 | Kctd13 | Kctd13 | magenta | 0.019326 |
| 228 | Echdc1 | Echdc1 | magenta | 0.019079 |
| 156 | Zfp418 | Zfp418 | magenta | 0.019033 |
| 72 | Ubxn2b | Ubxn2b | magenta | 0.019019 |
| 316 | Tmtc4 | Tmtc4 | magenta | 0.018454 |
| 236 | Cdk17 | Cdk17 | magenta | 0.017861 |
| 287 | Batf | Batf | magenta | 0.017261 |
| 20 | Ccdc181 | Ccdc181 | magenta | 0.01724 |
| 275 | Cant1 | Cant1 | magenta | 0.017207 |
| 19 | Gorab | Gorab | magenta | 0.017048 |
| 191 | Tspan32 | Tspan32 | magenta | 0.017028 |
| 144 | Shq1 | Shq1 | magenta | 0.016586 |
| 200 | Ano8 | Ano8 | magenta | 0.01641 |
| 292 | Phactr1 | Phactr1 | magenta | 0.016352 |
| 97 | Gm6745 | Gm6745 | magenta | 0.015544 |
| 279 | Laptm4a | Laptm4a | magenta | 0.015364 |
| 227 | Ccdc28a | Ccdc28a | magenta | 0.0138 |
| 76 | Gm12403 | Gm12403 | magenta | 0.012548 |
| 181 | Prkcb | Prkcb | magenta | 0.011681 |
| 341 | Lca5l | Lca5l | magenta | 0.011544 |
| 296 | Simc1 | Simc1 | magenta | 0.009489 |
| 41 | Ivd | Ivd | magenta | 0.008968 |
| 273 | Wipi1 | Wipi1 | magenta | 0.008348 |
| 379 | Nudt8 | Nudt8 | magenta | 0.008274 |
| 158 | Pvr | Pvr | magenta | 0.007831 |
| 396 | Npm3 | Npm3 | magenta | 0.007419 |

Table S3 Microglial interferon response module in human AD and MCI

| | ensgene | name | module | mm |
|---|---|---|---|---|
| 161 | CD163 | CD163 | salmon | 0.503175 |
| 125 | LY6E | LY6E | salmon | 0.463977 |
| 134 | IFITM3 | IFITM3 | salmon | 0.4289 |
| 5 | IFI6 | IFI6 | salmon | 0.410875 |
| 1 | ISG15 | ISG15 | salmon | 0.377123 |
| 8 | IFI44L | IFI44L | salmon | 0.354038 |
| 154 | IFIT1 | IFIT1 | salmon | 0.350797 |
| 153 | IFIT3 | IFIT3 | salmon | 0.340812 |
| 85 | F13A1 | F13A1 | salmon | 0.31491 |
| 162 | CLEC4E | CLEC4E | salmon | 0.312056 |
| 71 | SEPP1 | SEPP1 | salmon | 0.30934 |
| 165 | LYZ | LYZ | salmon | 0.296703 |
| 219 | BST2 | BST2 | salmon | 0.268431 |
| 79 | TGFBI | TGFBI | salmon | 0.265644 |
| 144 | MS4A4A | MS4A4A | salmon | 0.257309 |
| 233 | MX1 | MX1 | salmon | 0.247052 |
| 127 | ANXA1 | ANXA1 | salmon | 0.232329 |
| 9 | IFI44 | IFI44 | salmon | 0.213326 |
| 152 | IFIT2 | IFIT2 | salmon | 0.210522 |
| 26 | RSAD2 | RSAD2 | salmon | 0.209396 |
| 210 | LGALS3BP | LGALS3BP | salmon | 0.19831 |
| 138 | LYVE1 | LYVE1 | salmon | 0.195783 |
| 175 | EPSTI1 | EPSTI1 | salmon | 0.190855 |
| 14 | C1orf162 | C1orf162 | salmon | 0.182847 |
| 133 | IFITM1 | IFITM1 | salmon | 0.182164 |
| 131 | EGFL7 | EGFL7 | salmon | 0.178696 |
| 3 | RBP7 | RBP7 | salmon | 0.177585 |
| 16 | FCGR2B | FCGR2B | salmon | 0.176575 |
| 143 | MS4A6A | MS4A6A | salmon | 0.17574 |
| 193 | MT2A | MT2A | salmon | 0.170593 |
| 54 | CSTA | CSTA | salmon | 0.162992 |
| 170 | OAS1 | OAS1 | salmon | 0.162969 |
| 132 | IFITM2 | IFITM2 | salmon | 0.152799 |
| 25 | CMPK2 | CMPK2 | salmon | 0.151076 |
| 113 | AP1S2 | AP1S2 | salmon | 0.150932 |
| 174 | OASL | OASL | salmon | 0.148584 |
| 74 | IQGAP2 | IQGAP2 | salmon | 0.147221 |
| 31 | AC009501 | AC009501 | salmon | 0.144511 |
| 232 | MX2 | MX2 | salmon | 0.144123 |
| 93 | FAM26F | FAM26F | salmon | 0.141223 |
| 36 | POTEE | POTEE | salmon | 0.135909 |
| 171 | OAS3 | OAS3 | salmon | 0.131539 |
| 12 | DPYD | DPYD | salmon | 0.131295 |
| 145 | MS4A7 | MS4A7 | salmon | 0.126883 |

| | | | | |
|---|---|---|---|---|
| 182 | SNX6 | SNX6 | salmon | 0.12665 |
| 172 | OAS2 | OAS2 | salmon | 0.126469 |
| 213 | SIGLEC1 | SIGLEC1 | salmon | 0.125727 |
| 28 | EIF2AK2 | EIF2AK2 | salmon | 0.124208 |
| 45 | AZI2 | AZI2 | salmon | 0.119385 |
| 41 | STAT1 | STAT1 | salmon | 0.119127 |
| 18 | RABGAP1L | RABGAP1L | salmon | 0.115969 |
| 191 | EMP2 | EMP2 | salmon | 0.113304 |
| 180 | TNFSF13B | TNFSF13B | salmon | 0.113295 |
| 32 | VAMP5 | VAMP5 | salmon | 0.11027 |
| 109 | CLEC5A | CLEC5A | salmon | 0.106952 |
| 94 | ARHGAP18 | ARHGAP18 | salmon | 0.10611 |
| 92 | DSE | DSE | salmon | 0.105697 |
| 135 | IRF7 | IRF7 | salmon | 0.104136 |
| 104 | SAMD9L | SAMD9L | salmon | 0.103417 |
| 90 | FKBP1C | FKBP1C | salmon | 0.10263 |
| 49 | SHISA5 | SHISA5 | salmon | 0.101079 |
| 150 | MRC1 | MRC1 | salmon | 0.101026 |
| 168 | DRAM1 | DRAM1 | salmon | 0.099242 |
| 194 | MT1X | MT1X | salmon | 0.098557 |
| 229 | ODF3B | ODF3B | salmon | 0.098145 |
| 42 | SPATS2L | SPATS2L | salmon | 0.098098 |
| 98 | SNX10 | SNX10 | salmon | 0.097272 |
| 198 | ALOX15B | ALOX15B | salmon | 0.094841 |
| 58 | PLSCR1 | PLSCR1 | salmon | 0.093091 |
| 155 | IFIT5 | IFIT5 | salmon | 0.092994 |
| 55 | PARP9 | PARP9 | salmon | 0.088952 |
| 142 | MDK | MDK | salmon | 0.085214 |
| 226 | USP18 | USP18 | salmon | 0.083737 |
| 111 | PRKAG2-A | PRKAG2-A | salmon | 0.083726 |
| 76 | TNFAIP8 | TNFAIP8 | salmon | 0.083471 |
| 117 | NGFRAP1 | NGFRAP1 | salmon | 0.08294 |
| 7 | EPS15 | EPS15 | salmon | 0.081679 |
| 78 | CXCL14 | CXCL14 | salmon | 0.080875 |
| 116 | ARMCX1 | ARMCX1 | salmon | 0.08017 |
| 110 | GSTK1 | GSTK1 | salmon | 0.079385 |
| 39 | IFIH1 | IFIH1 | salmon | 0.079074 |
| 82 | SMIM3 | SMIM3 | salmon | 0.078848 |
| 66 | DDX60 | DDX60 | salmon | 0.078506 |
| 160 | RP5-940J5 | RP5-940J5 | salmon | 0.074874 |
| 149 | CASP4 | CASP4 | salmon | 0.074504 |
| 222 | ETHE1 | ETHE1 | salmon | 0.073677 |
| 114 | GPR82 | GPR82 | salmon | 0.071005 |
| 122 | FAM110B | FAM110B | salmon | 0.070946 |
| 225 | LILRB2 | LILRB2 | salmon | 0.069151 |
| 188 | TRIM69 | TRIM69 | salmon | 0.067185 |
| 120 | CLIC2 | CLIC2 | salmon | 0.063734 |

| | | | | |
|---|---|---|---|---|
| 139 | ZDHHC13 | ZDHHC13 | salmon | 0.063512 |
| 209 | GPRC5C | GPRC5C | salmon | 0.062794 |
| 34 | AC109826 | AC109826 | salmon | 0.061706 |
| 23 | FAM177B | FAM177B | salmon | 0.061148 |
| 163 | CLEC2B | CLEC2B | salmon | 0.060521 |
| 231 | IFNGR2 | IFNGR2 | salmon | 0.060045 |
| 60 | B3GALNT1 | B3GALNT1 | salmon | 0.059011 |
| 72 | GAPT | GAPT | salmon | 0.057694 |
| 11 | GBP1 | GBP1 | salmon | 0.057625 |
| 211 | RBFOX3 | RBFOX3 | salmon | 0.057421 |
| 183 | GCH1 | GCH1 | salmon | 0.057066 |
| 220 | ZNF429 | ZNF429 | salmon | 0.056794 |
| 106 | TRIP6 | TRIP6 | salmon | 0.056432 |
| 20 | IL10 | IL10 | salmon | 0.055874 |
| 59 | P2RY14 | P2RY14 | salmon | 0.055675 |
| 187 | LCMT2 | LCMT2 | salmon | 0.055229 |
| 73 | FCHO2 | FCHO2 | salmon | 0.054848 |
| 167 | PLXNC1 | PLXNC1 | salmon | 0.054204 |
| 27 | RP11-563N | RP11-563N | salmon | 0.053829 |
| 166 | DUSP6 | DUSP6 | salmon | 0.053338 |
| 123 | UBXN2B | UBXN2B | salmon | 0.052967 |
| 38 | KCNJ3 | KCNJ3 | salmon | 0.052522 |
| 96 | DYNLT1 | DYNLT1 | salmon | 0.051628 |
| 21 | LYPLAL1 | LYPLAL1 | salmon | 0.051622 |
| 146 | AHNAK | AHNAK | salmon | 0.051611 |
| 15 | CERS2 | CERS2 | salmon | 0.051348 |
| 157 | CUTC | CUTC | salmon | 0.050462 |
| 99 | TBRG4 | TBRG4 | salmon | 0.049837 |
| 47 | CCR5 | CCR5 | salmon | 0.049641 |
| 56 | DTX3L | DTX3L | salmon | 0.049401 |
| 81 | RBM22 | RBM22 | salmon | 0.048242 |
| 86 | ABT1 | ABT1 | salmon | 0.048084 |
| 205 | CALCOCO2 | CALCOCO2 | salmon | 0.048012 |
| 91 | SESN1 | SESN1 | salmon | 0.047604 |
| 202 | PLXDC1 | PLXDC1 | salmon | 0.04723 |
| 230 | U2AF1L5 | U2AF1L5 | salmon | 0.0471 |
| 186 | TRMT61A | TRMT61A | salmon | 0.047017 |
| 61 | FXR1 | FXR1 | salmon | 0.046916 |
| 95 | SYNE1 | SYNE1 | salmon | 0.046215 |
| 48 | ELP6 | ELP6 | salmon | 0.045865 |
| 192 | EARS2 | EARS2 | salmon | 0.04572 |
| 67 | SAP30 | SAP30 | salmon | 0.045693 |
| 115 | WDR45 | WDR45 | salmon | 0.045508 |
| 37 | KYNU | KYNU | salmon | 0.04529 |
| 10 | PRKACB | PRKACB | salmon | 0.044633 |
| 68 | SCRG1 | SCRG1 | salmon | 0.044397 |
| 129 | MRPL50 | MRPL50 | salmon | 0.043695 |

| | | | | |
|---|---|---|---|---|
| 51 | C3orf38 | C3orf38 | salmon | 0.043403 |
| 65 | ZNF330 | ZNF330 | salmon | 0.042861 |
| 200 | LGALS9C | LGALS9C | salmon | 0.042651 |
| 179 | TEX30 | TEX30 | salmon | 0.042217 |
| 189 | C15orf61 | C15orf61 | salmon | 0.04211 |
| 119 | DNASE1L1 | DNASE1L1 | salmon | 0.04209 |
| 148 | INTS4 | INTS4 | salmon | 0.042038 |
| 40 | MTX2 | MTX2 | salmon | 0.041482 |
| 77 | C5orf56 | C5orf56 | salmon | 0.041301 |
| 158 | SFR1 | SFR1 | salmon | 0.041121 |
| 156 | PCGF5 | PCGF5 | salmon | 0.040804 |
| 184 | MAX | MAX | salmon | 0.040774 |
| 128 | STX17 | STX17 | salmon | 0.040671 |
| 17 | SUCO | SUCO | salmon | 0.040621 |
| 62 | RTP4 | RTP4 | salmon | 0.040573 |
| 75 | RHOBTB3 | RHOBTB3 | salmon | 0.040551 |
| 69 | GPM6A | GPM6A | salmon | 0.040502 |
| 80 | SLC35A4 | SLC35A4 | salmon | 0.040441 |
| 44 | HACL1 | HACL1 | salmon | 0.040401 |
| 87 | TAP1 | TAP1 | salmon | 0.040237 |
| 84 | DUSP22 | DUSP22 | salmon | 0.039892 |
| 35 | SLC35F5 | SLC35F5 | salmon | 0.039805 |
| 141 | IMMP1L | IMMP1L | salmon | 0.039674 |
| 101 | TMEM60 | TMEM60 | salmon | 0.039485 |
| 108 | ALKBH4 | ALKBH4 | salmon | 0.039137 |
| 2 | TNFRSF14 | TNFRSF14 | salmon | 0.039131 |
| 112 | LINC01003 | LINC01003 | salmon | 0.03905 |
| 13 | PRMT6 | PRMT6 | salmon | 0.038928 |
| 206 | ABCC3 | ABCC3 | salmon | 0.0389 |
| 103 | FZD1 | FZD1 | salmon | 0.038662 |
| 199 | TMEM220 | TMEM220 | salmon | 0.038227 |
| 216 | TPGS1 | TPGS1 | salmon | 0.038023 |
| 207 | AKAP1 | AKAP1 | salmon | 0.037914 |
| 50 | PPP4R2 | PPP4R2 | salmon | 0.037833 |
| 24 | OPN3 | OPN3 | salmon | 0.037628 |
| 212 | TIMM21 | TIMM21 | salmon | 0.037586 |
| 214 | SLC35C2 | SLC35C2 | salmon | 0.037497 |
| 63 | NIPAL1 | NIPAL1 | salmon | 0.037152 |
| 224 | NAPSA | NAPSA | salmon | 0.037061 |
| 57 | ACAD9 | ACAD9 | salmon | 0.035852 |
| 178 | DIS3 | DIS3 | salmon | 0.03576 |
| 201 | SLFN5 | SLFN5 | salmon | 0.03497 |
| 181 | AP4S1 | AP4S1 | salmon | 0.03497 |
| 218 | KEAP1 | KEAP1 | salmon | 0.034894 |
| 88 | RXRB | RXRB | salmon | 0.03441 |
| 30 | FOXN2 | FOXN2 | salmon | 0.034219 |
| 19 | STX6 | STX6 | salmon | 0.033805 |

| | | | | |
|---|---|---|---|---|
| 227 | APOL1 | APOL1 | salmon | 0.033799 |
| 100 | STYXL1 | STYXL1 | salmon | 0.033092 |
| 173 | C12orf49 | C12orf49 | salmon | 0.033066 |
| 137 | RNF141 | RNF141 | salmon | 0.032963 |
| 204 | NSF | NSF | salmon | 0.032737 |
| 217 | CTD-3116E | CTD-3116E | salmon | 0.03272 |
| 140 | ARL14EP | ARL14EP | salmon | 0.032455 |
| 223 | GEMIN7 | GEMIN7 | salmon | 0.031905 |
| 196 | PITPNA-AS | PITPNA-AS | salmon | 0.031494 |
| 52 | IQCB1 | IQCB1 | salmon | 0.03142 |
| 46 | LRRFIP2 | LRRFIP2 | salmon | 0.03118 |
| 102 | CROT | CROT | salmon | 0.031156 |
| 151 | ACBD5 | ACBD5 | salmon | 0.031111 |
| 203 | STARD3 | STARD3 | salmon | 0.030657 |
| 121 | SH2D4A | SH2D4A | salmon | 0.030294 |
| 164 | CCDC91 | CCDC91 | salmon | 0.029685 |
| 215 | PRPF6 | PRPF6 | salmon | 0.029461 |
| 22 | IARS2 | IARS2 | salmon | 0.029177 |
| 228 | PICK1 | PICK1 | salmon | 0.02911 |
| 105 | PPP1R35 | PPP1R35 | salmon | 0.028691 |
| 118 | RNF113A | RNF113A | salmon | 0.028108 |
| 33 | KRCC1 | KRCC1 | salmon | 0.027684 |
| 107 | SH2B2 | SH2B2 | salmon | 0.027121 |
| 130 | MAPKAP1 | MAPKAP1 | salmon | 0.027097 |
| 147 | RELA | RELA | salmon | 0.027033 |
| 6 | KHDRBS1 | KHDRBS1 | salmon | 0.027032 |
| 97 | ANKMY2 | ANKMY2 | salmon | 0.02691 |
| 124 | TMEM65 | TMEM65 | salmon | 0.026825 |
| 197 | TMEM102 | TMEM102 | salmon | 0.026324 |
| 43 | MKRN2 | MKRN2 | salmon | 0.026146 |
| 177 | DHRS12 | DHRS12 | salmon | 0.02562 |
| 89 | CD2AP | CD2AP | salmon | 0.025477 |
| 190 | ZNF200 | ZNF200 | salmon | 0.025009 |
| 221 | CEACAM4 | CEACAM4 | salmon | 0.02398 |
| 64 | NUP54 | NUP54 | salmon | 0.023443 |
| 70 | UFSP2 | UFSP2 | salmon | 0.023097 |
| 29 | MAP4K3 | MAP4K3 | salmon | 0.022506 |
| 126 | FAM122A | FAM122A | salmon | 0.022451 |
| 83 | BOD1 | BOD1 | salmon | 0.022145 |
| 136 | PNPLA2 | PNPLA2 | salmon | 0.021843 |
| 169 | MTERF2 | MTERF2 | salmon | 0.021568 |
| 176 | PHF11 | PHF11 | salmon | 0.021565 |
| 159 | STK32C | STK32C | salmon | 0.021263 |
| 208 | PPM1D | PPM1D | salmon | 0.021169 |
| 53 | EAF2 | EAF2 | salmon | 0.020776 |
| 195 | CENPBD1 | CENPBD1 | salmon | 0.020483 |
| 4 | MAP3K6 | MAP3K6 | salmon | 0.019881 |

| | | | | |
|---|---|---|---|---|
| 185 | WDR20 | WDR20 | salmon | 0.017543 |
